# Long-read genome sequencing accelerated the cloning of *Pm69* by resolving the complexity of a rapidly evolving resistance gene cluster in wheat

**DOI:** 10.1101/2022.10.14.512294

**Authors:** Yinghui Li, Zhen-Zhen Wei, Hanan Sela, Liubov Govta, Valentyna Klymiuk, Rajib Roychowdhury, Harmeet Singh Chawla, Jennifer Ens, Krystalee Wiebe, Valeria Bocharova, Roi Ben-David, Prerna B. Pawar, Samidha Jaiwar, István Molnár, Jaroslav Doležel, Curtis J. Pozniak, Tzion Fahima

## Abstract

Gene cloning in repeat-rich polyploid genomes remains challenging. Here we describe a strategy for overcoming major bottlenecks in the cloning of the powdery mildew (Pm) resistance gene (R-gene) *Pm69* derived from tetraploid wild emmer wheat (WEW). A conventional positional cloning approach encountered suppressed recombination due to structural variations, while chromosome sorting yielded an insufficient purity level. A *Pm69* physical map, constructed by assembling ONT long-read genome sequences, revealed a rapidly evolving nucleotide-binding leucine-rich repeat (NLR) R-gene cluster. A single candidate NLR was identified within this cluster by anchoring RNASeq reads of susceptible mutants to ONT contigs and was validated by the virus-induced gene silencing (VIGS) approach. *Pm69*, comprising Rx_N with RanGAP interaction sites, NB-ARC, and LRR domains, is probably a newly evolved NLR discovered only in one location across the WEW distribution range in the Fertile Crescent. *Pm69* was successfully introgressed into durum and bread wheat, and a diagnostic molecular marker could be used to accelerate its deployment and pyramiding with other resistance genes.

## Introduction

Powdery mildew (Pm) caused by the biotrophic fungus *Blumeria graminis* f. sp. *tritici* (*Bgt*) is one of the most destructive wheat diseases worldwide. To date, more than 110 *Pm* resistance genes/alleles have been identified and mapped in wheat and its wild relatives^1^, of which only about 10% have been cloned. *Pm3b/Pm8, Pm2a, Pm21, Pm60/MlIW172, Pm5e, Pm41* and *Pm1a* encode nucleotide-binding leucine-rich repeat (NLR) proteins^2–10^. *Pm24* encodes a tandem kinase protein and *Pm4* encodes a putative chimeric protein of a serine/threonine kinase, multiple C2 domains, and transmembrane regions^11,12^. *Pm38 (Lr34/Yr18/Sr57*, ABC transporter) and *Pm46 (Lr67/Yr46/Sr55*, hexose transporter) show broad-spectrum adult plant resistance to powdery mildew and rust diseases^13,14^. However, *Pm* resistance genes are frequently overcome due to the rapid evolution of *Bgt* isolates^15^. Thus, the identification and cloning of novel disease resistance genes (R-genes) and deployment into cultivated wheat germplasm can enrich the repertoire of genes available for resistance breeding.

Positional cloning has been widely used to clone genes. The published reference genomes of several wheat species are serving as powerful tools for the dissection of the target genomic regions and provide a reliable source for candidate gene prediction^16^. However, genes responsible for the phenotype of interest may be absent from the reference genotypes, especially when the target genes are derived from wild relatives^17^. Analysis of 16 hexaploid wheat genomes revealed that only 31–34% of the NLR signatures are shared across all the genomes, indicating that this gene family is subjected to rapid evolutionary processes^16,18^. Analysis of the Chinese Spring (CS) bread wheat genome assembly revealed 2,151 NLR-like genes, of which 1,298 were arranged in 547 gene clusters, many of them showing more than 75% similarity within each cluster, likely formed by tandem duplications^19^. Such clusters of highly similar genes make it particularly challenging to localize a causal R-gene, especially when relying heavily on published reference genomes. The construction of a Bacterial Artificial Chromosome (BAC) library with 100-200 kb insert size may be useful for assembling high-quality physical maps to support positional cloning^20^, however, this technology is time and labor-intensive, especially when working with complex genomes, such as wheat.

To overcome the limitations of R-genes’ positional cloning and reduce the genomic complexity of wheat, several methods based on next-generation sequencing (NGS) have been developed and their efficiency was demonstrated for cloning of novel R-genes^21^. These methods include Mutagenesis and Resistance gene Enrichment and Sequencing (MutRenSeq)^22^, Association genetics with R gene enrichment Sequencing (AgRenSeq)^23^, Mutant Chromosome flow sorting and short-read Sequencing (MutChromSeq)^3^, and Targeted Chromosome-based Cloning via long-range Assembly (TACCA)^24^. However, these methods still have some limitations. For example, MutRenSeq and AgRenSeq only identify NLR genes that can be captured by hybridization, while TACCA and MutChromSeq rely on the purification of individual chromosomes that carry mutations in the target genes^21,25^. Moreover, Bulked segregant RNA sequencing (BSR-Seq) and bulked segregant Core Genome Targeted sequencing (CGT-Seq) have been used successfully for identifying R-genes, but these methods rely on the selection scales of individuals for mixing genomic bulked pools^6,26^.

Wheat has a large and complex genome with more than 85% repetitive sequences making genome sequence assembly challenging^27^, particularly when using short-read (100-250 bp) sequencing technologies. With the rapid development of sequencing methodologies, the long-read sequencing platforms such as Pacific Biosciences (PacBio) single-molecule real-time (SMRT) sequencing and Oxford Nanopore Technologies (ONT) sequencing generate sufficiently long (>10 kb) reads that result in more contiguous sequence assemblies^28,29^. Long-read sequencing technologies have been successfully used for barley and bread wheat genome assemblies^30–33^. These technologies can improve physical mapping certainty, detection of structural variants, and transcript isoform identification^16,34,35^.

Wild emmer wheat (*Triticum turgidum* ssp. *dicoccoides*, WEW, *2n = 4x =* 28, AABB), the tetraploid progenitor of hexaploid bread wheat (*T. aestivum*, 2n=6x=42, AABBDD), is a valuable genetic resource for resistance genes. More than 20 *Pm* resistance genes have been identified and mapped in WEW^36^. Among them, only two were cloned, the *Pm41* localized on chromosome arm 3BL and *TdPm60* localized on chromosome arm 7AL. *Pm41* and *TdPm60* showed abundance of 1.81% and 25.6% among the tested natural WEW accessions, respectively^8,9,37^. *Pm69 (PmG3M*), identified from WEW accession G305-3M, is a dominant gene conferring a wide-spectrum resistance to *Bgt* isolates from around the globe^38^. *Pm69* is the only gene derived from WEW that was mapped to the telomeric region of chromosome arm 6BL^36,38,39^.

In the current study, we have developed a high-resolution genetic map of chromosome arm 6BL that contains *Pm69*, however, we encountered suppressed recombination due to structural variations within the target region and therefore, conventional positional cloning was impossible. To resolve the complexity of this genomic region, we used the long-read Oxford Nanopore Technology (ONT) for whole genome sequencing of the *Pm69* donor line (G305-3M) coupled with short-read transcriptome sequencing (RNA-seq) of *pm69* susceptible mutants. A single candidate NLR was identified within a rapidly evolving NLR gene cluster and validated using virus-induced gene silencing (VIGS). Although *Pm69* is providing resistance to numerous *Bgt* isolates, it was found to be a very rare allele among WEW natural populations, indicating that it may represent a newly evolved NLR within a rapidly evolving cluster. We introgressed *Pm69* into cultivated wheat and pyramided it with three yellow rust (caused by the fungus *Puccinia striiformis* f.sp. *tritici, Pst*) resistance genes (*Yr5, Yr15*, and *Yr24*), to provide a valuable resource for disease resistance breeding in wheat.

## Results

### The challenges of chromosome walking in a complex structurally variable region

A recombinant inbred line (RIL) mapping population segregating for *Pm69* was generated by crossing the susceptible durum wheat (*Triticum turgidum* ssp. *durum*) Langdon (LDN) with the resistant WEW accession G305-3M. G305-3M and the F1 generation (G305-3M × LDN) showed post-haustorial immune responses to *Bgt* #70 accompanied by intracellular ROS production and host cell death, while LDN showed a highly susceptible response resulting in the development of massive fungal pathogen colonies (Figure 1a). G305-3M showed a wide-spectrum resistance against 55 tested *Bgt* isolates, originating from four continents Asia, Europe, North America, and South America (Table S1). For fine mapping of *Pm69*, we screened 5500 F2 plants (11,000 gametes) with DNA markers *uhwk386* and *uhwk399* to develop 147 F4-7 RILs that carried informative recombination events within the target region. In total, additional 33 DNA markers were developed and mapped within this genetic region (Figure S1, Table S2), enabling to map *Pm69* within a 0.21 cM interval between markers *uhw367* and *uhwk389* (Figure 1, Table S3).

**Figure 1.**
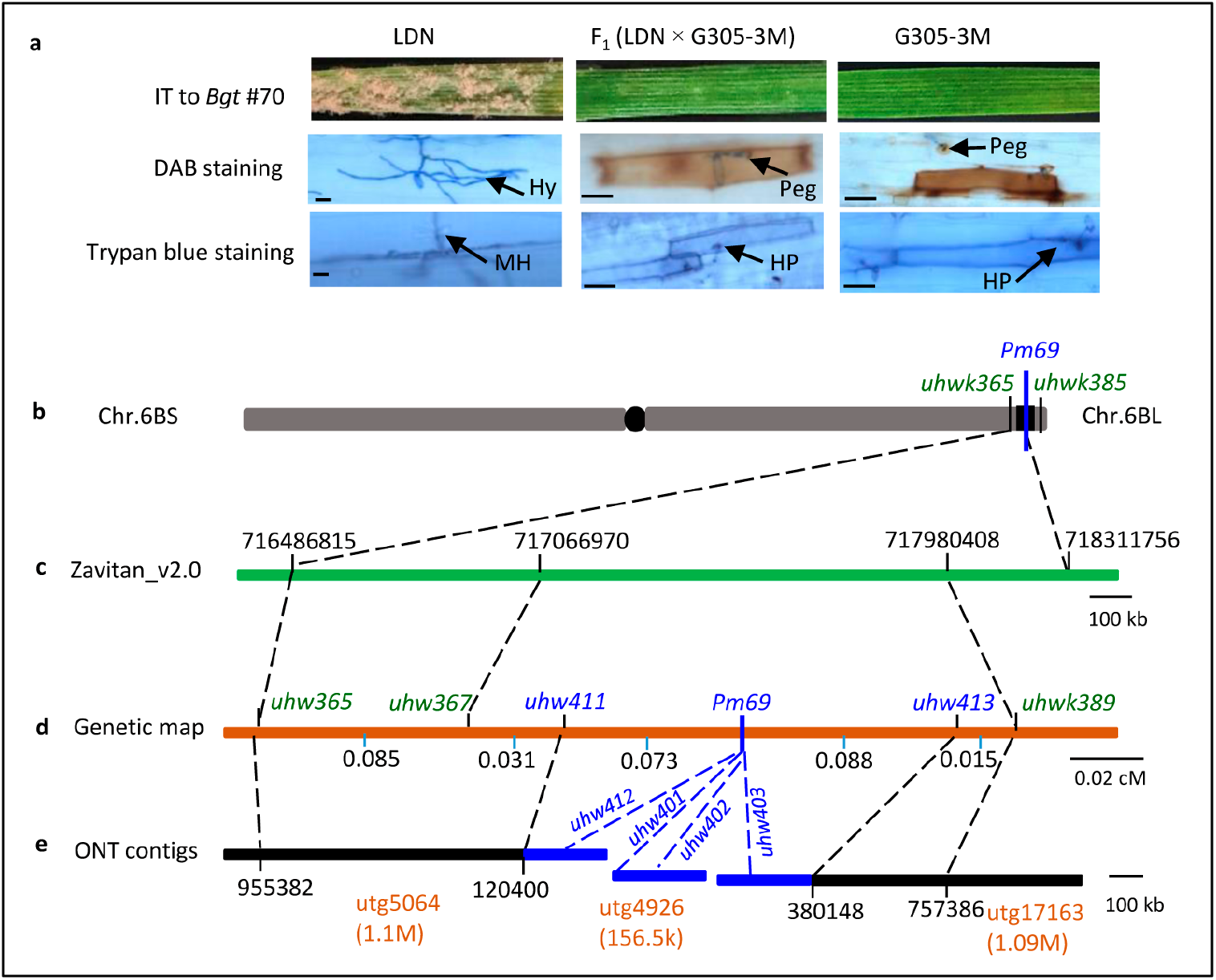
Fine mapping of *Pm69* from WEW G305-3M. **a** Macroscopic observation of the response of G305-3M, LDN and F1 leaves to *Bgt* # 70 infection at 7 days post-infection (dpi); DAB staining of leaves at 3 dpi to detect ROS accumulation visualized as Reddish-brown coloration; Trypan blue staining of leaves at 3 dpi to visualize fungal structures and plant cell death as blue coloration; MH: mature haustorium; HP: haustorial primordium; Hy: hyphae. Scale bars = 50 μm. **b** The *Pm69* genetic region on the long arm of wheat chromosome 6B. **c** The locations of *Pm69* flanking genetic markers on the WEW _v2.0 reference genome. **d** The genetic map of *Pm69*. The green-colored markers were developed based on WEW _v2.0, the blue-colored markers were developed based on ONT contigs. **e** The 6BL ONT contig assembly in the *Pm69* genetic region of G305-3M. The physical map of the *Pm69* region is marked in blue.

We anchored these closely linked genetic markers to three wheat reference genomes and found that *Pm69* genetic map showed a higher collinearity level with the WEW_v2.0 assembly than with that of durum wheat Svevo (RefSeq Rel. 1.0) and bread wheat Chinese Spring (IWGSC RefSeq v1.0) (Figure S2). As Zavitan, Svevo and Chinese Spring are susceptible to *Bgt* #70 (Figure S3), we did not expect to find their functional *pm69* alleles. In Zavitan, this region is spanning 1.02 Mb comprising 23 genes, of which 18 are NLRs (Table S4). However, 10 markers physically mapped in this region showed co-segregation with the *Pm69* phenotype, suggesting suppressed recombination (Figure S2). In addition, 12 markers developed based on these candidate genes amplified PCR products in the susceptible parent Langdon, but not in the *Pm69* donor line G305-3M (Table S5). Taken together, these results indicated that structural variation between G305-3M and LDN probably caused the suppression of recombination around *Pm69* genetic region and thus prevented further chromosome walking towards *Pm69* based on the published reference genomes.

### The challenges of using MutChromSeq based on chromosome sorting in tetraploid wheat

As an alternative approach, we attempted to use MutChromSeq for cloning of *Pm69^3^*. To produce *pm69* loss-of-function mutants, we used ethyl methane sulfonate (EMS) treatment to mutagenize the WEW accession G305-3M and resistant F7-8 RILs. A total of five independent susceptible mutant lines (four in the G305-3M background and one in the resistant RIL background) were detected and grown to M4 generation (Figure S4).

We then attempted to isolate 6B chromosomes from G305-3M and LDN by flow cytometry-based chromosome sorting. Because of similar size and GAA microsatellite content, chromosomes 6B, 1B, 7B, 4B and 5B formed a composite and poorly resolved population on bivariate flow karyotype (Figure S5). We sorted chromosomes from the composites in hopes of further enriching chromosome 6B, but were only able to achieve a maximum purity of 47% and 51% in the case of G305-3M and LDN, respectively (Figure S5). The sorted fractions were contaminated by other chromosomes (1B, 7B, 4B and 5B in case of G305-3M, and chromosomes 1B, 7B and 5B in case of LDN) (Figure S5).

We performed an additional attempt to flow sort chromosome 6B from the hexaploid introgression line SC28RRR-26 (a bread wheat introgression line cv. Ruta + *Pm69*). The position of chromosome 6B population on the flow karyotype indicated its higher DNA content relative to other B-genome chromosomes. Improved discrimination of chromosome 6B resulted in a higher purity (73.1%) of the fraction flow sorted from the hexaploid wheat background (Figure S6). At that moment EMS-derived mutants of the introgression line were not available yet and it would take at least a couple of years to create these mutants, thus, alternative approaches were considered.

### Dissection of the structural variation complexity by ONT sequencing of G305-3M

The ONT long-read sequencing technology offered an attractive alternative to rapidly generate long contigs that span the *Pm69* genomic interval, overcoming difficulties conferred by the structural variations in the target gene region. ONT sequencing was used for whole-genome sequencing (~23x coverage) of WEW accession G305-3M. After assembling the reads, we obtained 2,489 Contigs, with N50 value of 11.2 Mb and N90 value of 2.6 Mb (Figure S7). The longest contig was 70.65 Mb, and the total length of the genome assembly was 10 GB (Table S6), which is typical for the tetraploid wheat genome^40,41^.

### Construction of *Pm69* physical map using ONT contigs

To construct the *Pm69* physical map, we anchored the ONT contigs to the *Pm69* genetic map using the co-dominant PCR markers. Two contigs utg17163 (1.1 Mb) and utg5064 (1.09 Mb) were perfectly anchored to the genetic map by four *Pm69* flanking markers (Table S7). An additional ONT contig utg4926 (156.5 kb) was identified by searching for gene sequences that reside in the *Pm69* collinear region of the WEW_v2.0 genome (Table S7). Based on the three identified ONT contigs, we have developed 15 markers that were incorporated into the genetic map by graphical genotyping of the RIL population (Table S8). Finally, the *Pm69* co-segregating markers spanned part of contig utg5064 (0-120400 bp), the whole contig utg4926 (0-156528 bp), and part of contigs utg17163 (0-380148 bp) (Figure 1, Table S8).

A comparison of G305-3M ONT contigs with the 6B pseudomolecules of WEW_ v2.0 and the Svevo reference genomes showed a very low level of collinearity around the *Pm69* genetic region, indicating massive structural rearrangements between them (Figure 2a and 2b, respectively). In contrast, more distant regions flanking *Pm69* showed a high collinearity level between the G305-3M contigs and the two reference genomes (Figure 2a and 2b).

**Figure 2.**
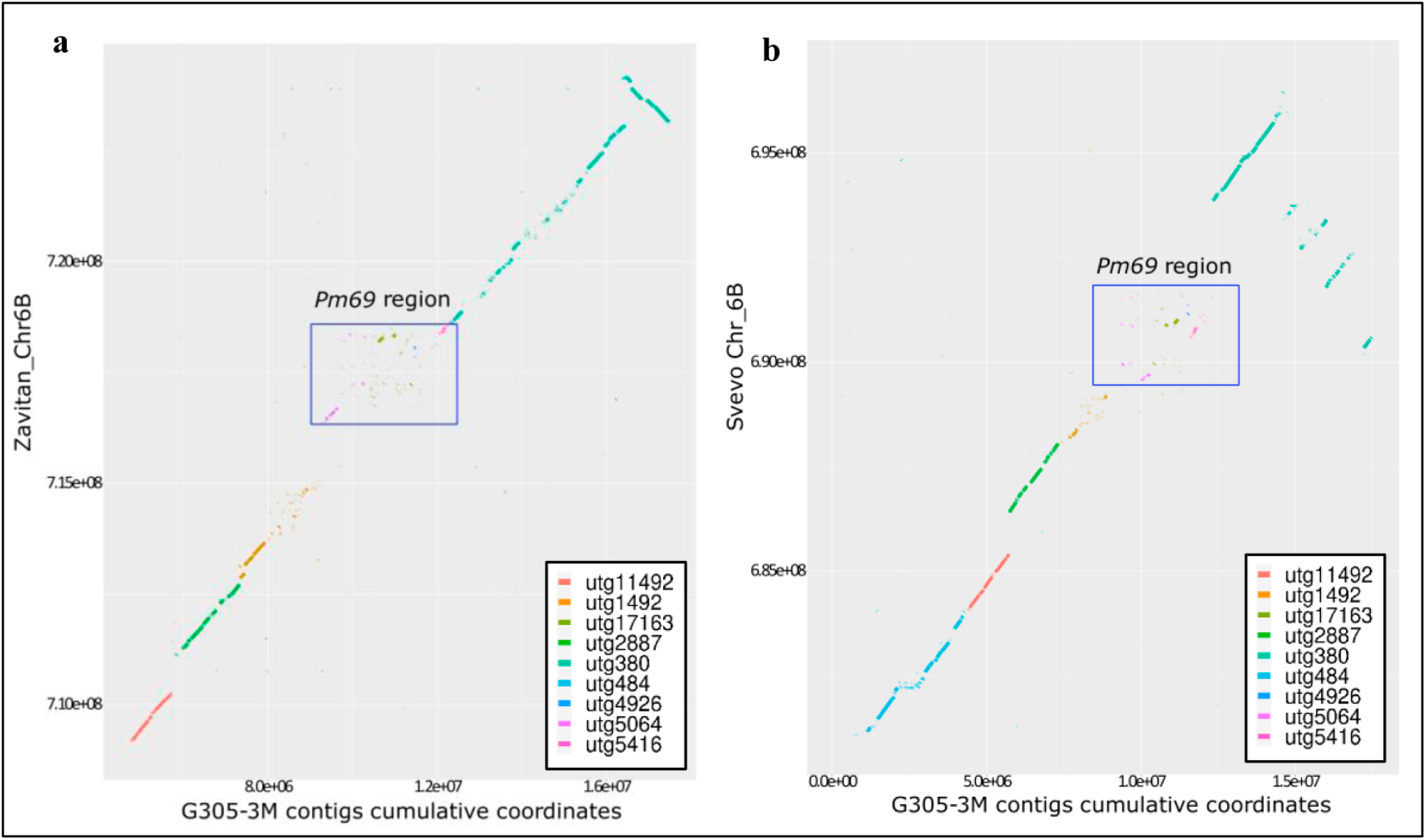
Comparisons of the G305-3M ONT contigs with the 6B pseudomolecule of WEW_v2.0 (a) and durum wheat Svevo RefSeq Rel. 1.0 (b) around the *Pm69* genetic region. X-axis: the cumulative length of G305-3M contigs; Y-axis: the physical location of 6B pseudomolecule in the reference genome. Contigs utg380-utg17163 marked with different colors belong to the ONT assembly of G305-3M.

### Identification of *Pm69* candidate gene by ONT-MutRNAseq

To identify a candidate for *Pm69* in the final physical interval, we mapped the RNAseq reads of four independent EMS-derived susceptible mutants onto the G305-3M ONT contigs (Figure 3a, 3b, S8, and Table S9). Using this approach, we revealed eight expressed candidate genes in the *Pm69* physical region, seven in utg17163, one in utg5046, and none in utg4926. Seven candidates were predicted as NLRs and one as a FAR1 Related Sequence (FRS) transcription factor (Figure 3b, Table S10). Only one gene spanning 12280 bp on G305-3M contig utg17163, named *NLR-6*, had five different point mutations in the four susceptible mutants (Figure S8, Table S11). These five mutations in *NLR-6* were all G/C to A/T transitions typical of EMS mutagenesis, while the remaining genes in this region did not have any mutations.

**Figure 3.**
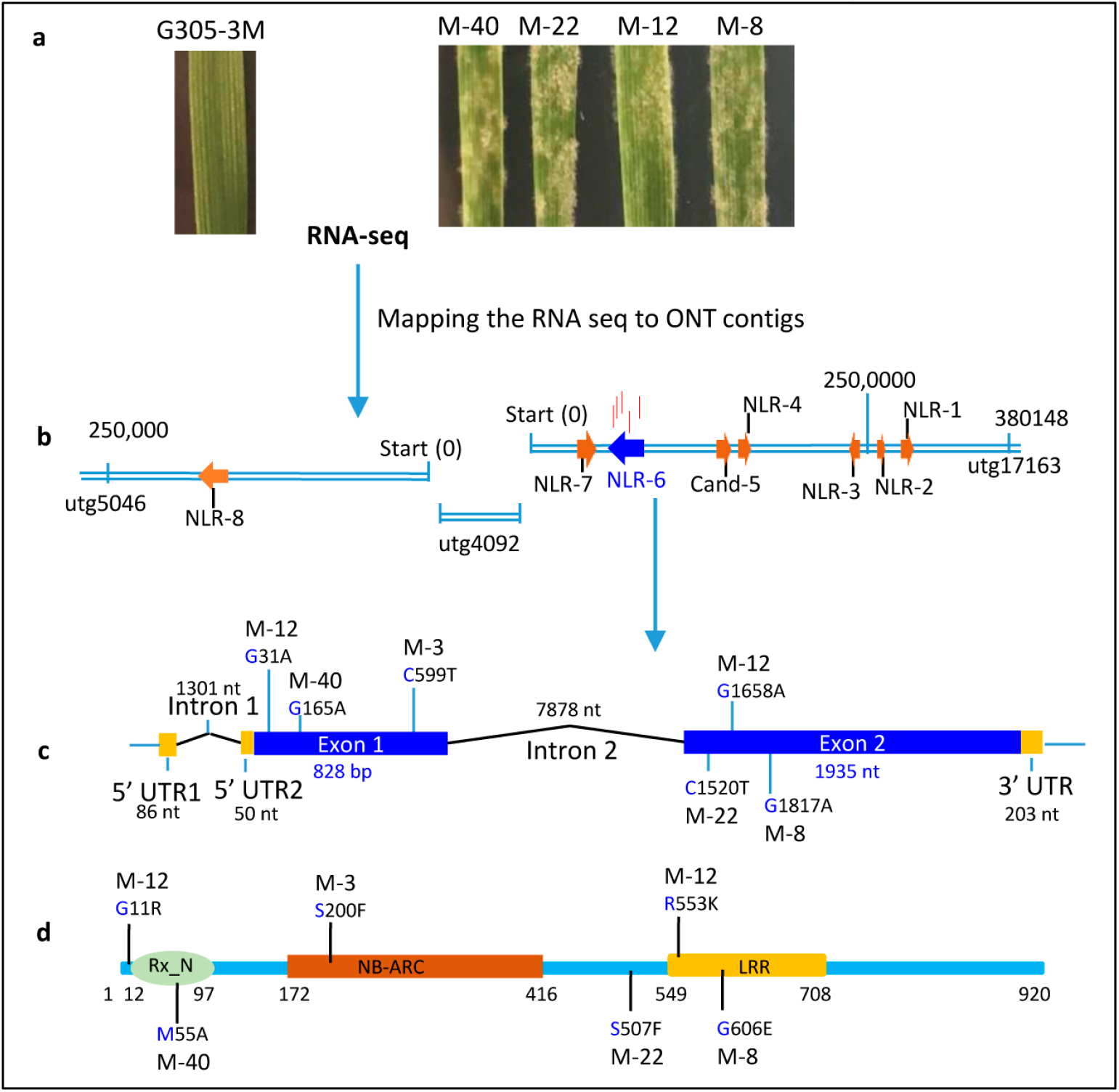
The workflow of identification of *Pm69* on the ONT contigs by MutRNAseq. **a** The phenotypes of EMS-derived mutants (M-40, M-22, M-12 and M-18) and wild type (G305-3M) infected with *Bgt* #70. M-40, M-22, M-12 and M-18 and G305-3M were analyzed by RNA-seq to identify SNPs among the three ONT contigs. **b** The expressed genes in the *Pm69* genetic region identified by aligning the RNAseq reads to G305-3M ONT contigs. Vertical red lines: the locations of SNPs in the susceptible mutants identified after mapping the RNASeq reads to G305-3M ONT contigs. **c** The distribution of the independent mutations. **d** The predicted structure of Pm69 protein with Rx_N, NB-ARC and LRR domains, and the precise location of all the identified mutants.

The predicted 2763 bp coding sequence (CDS) structure of *NLR-6* was validated by Sanger sequencing using G305-3M cDNA. The ~12.3 kb genomic region of the gene contains two introns, one of them located in the 5’ UTR region (Figure 3c). All the identified point mutations of *NLR-6* were validated by Sanger sequencing and were confirmed as missense mutations. Moreover, the resistant sister lines, derived from the same M0 plants, harbored the *Pm69* wild-type allele. An additional susceptible mutant line (M-3), identified by Sanger sequencing, also contained a point mutation (C599T, protein S200F) in NLR-6 (Figure 3c, Table S11). The predicted structure of the NLR-6 protein contains an Rx_N domain with RanGAP interaction sites, NB-ARC domain, and LRR domain (Figure 3d).

The expression pattern of *Pm69 (NLR-6*) was further examined at different time points from 0 to 72 hours post-infection (hpi) in both non-inoculated (mock) and inoculated G305-3M plants. The expression level of *Pm69* did not change significantly in the *Bgt* #70-infected plants during 0-16 hpi, compared with the non-inoculated control. However, *Pm69* expression was significantly decreased (p < 0.05) in the inoculated plants at 16-72 hpi (Figure S9).

### Functional validation of *Pm69* by virus-induced gene silencing (VIGS)

Virus-induced gene silencing (VIGS) constructs were designed to target two genomic positions in *Pm69* (Figure S10). We used a phytoene desaturase (*PDS*) gene silencing constructs to test the efficacy of the VIGS system in tetraploid (G305-3M) and hexaploid (a bread wheat introgression line cv. Ruta + *Pm69*) backgrounds. The inoculation with the *Pm69*-silencing constructs resulted in susceptibility to *Bgt* #70 of the 4^th^-5^th^ leaves in G305-3M and *Pm69* introgression line, while the negative controls showed no visible *Bgt* symptoms on the leaves (Figure 4a and 4b). Histopathological characterization showed that *Pm69* control cells (BSMV:GFP) accumulated intracellular ROS and prevented the invasion of the *Bgt* germinating spores. Silencing of PDS is visualized as white streaks resulting from photobleached chlorophyll^42^ (Figure 4a), while in the *Pm69* VIGS silenced leaves, we detected a mosaic pattern of germinating spores that invaded the cells successfully and developed colonies, alongside spores that activated HR cell death responses, as in the control resistant plants (Figure S11). Quantitative reverse transcription PCR (qRT–PCR) showed a significant reduction of expression levels of *Pm69* mRNA in the *Pm69* VIGS silenced leaves compared with GFP silenced leaves (p < 0.05; Figure 4c and S11). Silencing of *Pm69* in G305-3M and the introgression line (Ruta + *Pm69*) resulted in susceptible phenotypes, therefore, providing functional validation for the role of the *Pm69* gene (NLR-6) in conferring resistance against wheat powdery mildew.

**Figure 4.**
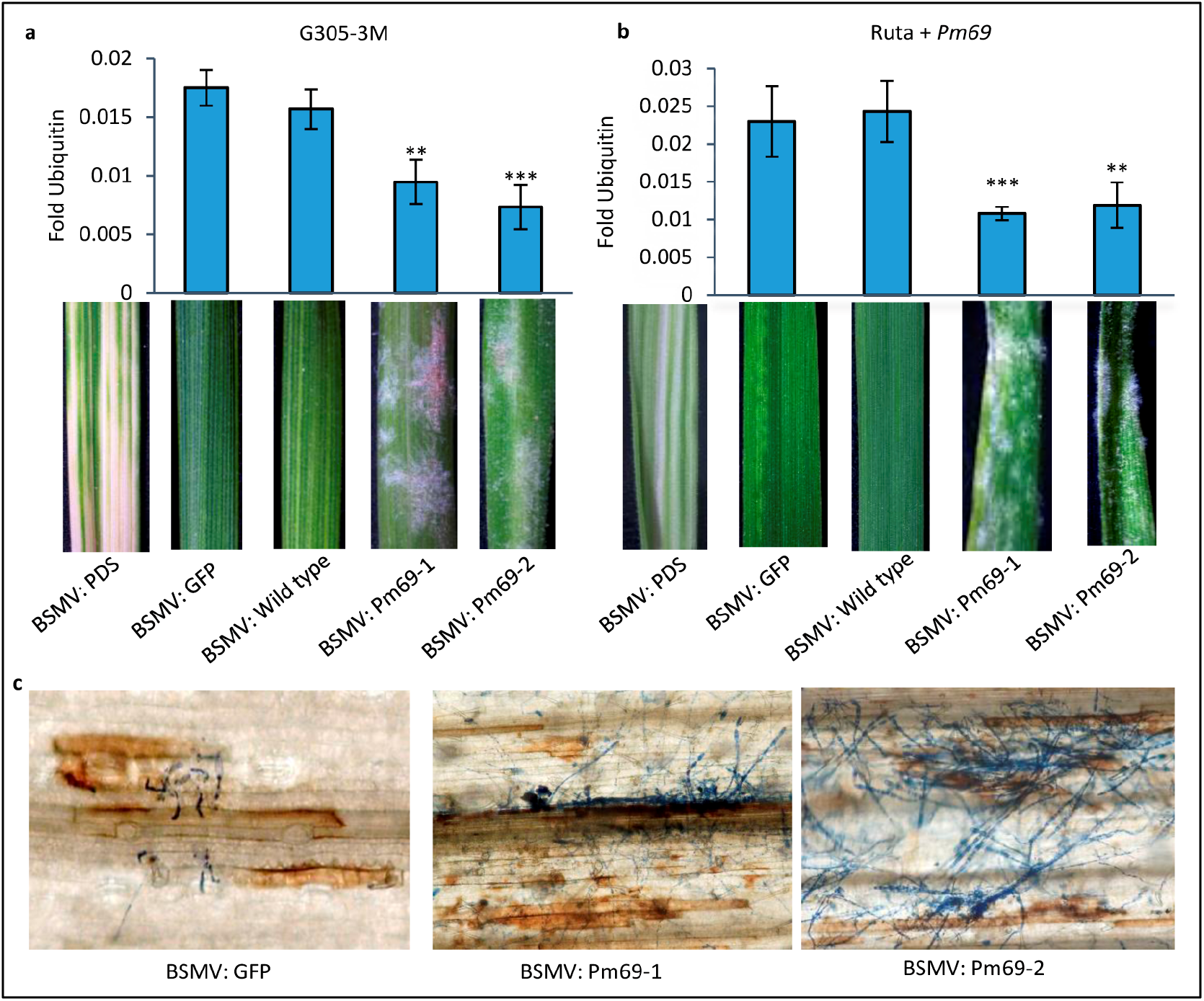
Functional validation of *Pm69* candidate gene through VIGS. **a** VIGS of *Pm69* candidate gene in G305-3M. **b** VIGS of *Pm69* candidate gene in the introgression line (Ruta+*Pm69*). BSMV:GFP was used as a negative control. BSMV:wild type, the wild type of *pCa-BSMV-γ* vector, was used as a negative control. BSMV:PDS is the PDS gene silencing construct used to check the efficiency of the VIGS system; BMSV:Pm69-1 and Pm69-2 were the constructs that targeted 5’UTR-Rx_N domain and NB-ARC domain, respectively. Asterisks indicate the level of significance by t-test of the *Pm69* expression levels in the target plants compared with the negative control of BSMV: wild type, p < 0.05 (*), p < 0.01 (**), p < 0.001 (***). **c** Histopathology characterization of the *Pm69* wild-type plants (BSMV:GFP) and *Pm69*-silencing plants (BSMV:Pm69-1 and BSMV:Pm69-2). DAB and Coomassie brilliant blue staining of *Pm69* or *GFP* silencing leaves in G305-3M at 7 dpi with *Bgt* #70.

### The *Pm69* is a rare allele

A search in the published wheat genome sequences identified several *Pm69* homologs/orthologs on chromosomes 6B and 6D from tetraploid and hexaploid wheat, as well as on chromosome 6 of *Aegilops tauschii*, which showed 87.55-88.77% protein sequence similarity with Pm69 (Table S12, Figure S12). Most of the closest NLR homologs from 6A chromosomes showed less than 50% protein similarity with Pm69, except for the *T. aestivum* cv. Julius 6A NLR that showed 85.46% similarity (Table S12). Protein sequence analysis of Pm69 homologs showed that most of the diversity was present in the LRR domain (Figure S13). Phylogenetic analysis showed that 24 homologs that showed 85-89% similarity to Pm69 were separated into several clades, most of them containing 2-7 identical proteins (Figure S12). Phenotyping of 11 representative accessions that cover the diversity among these 24 homologs (marked in orange in Figure S12) showed that all were susceptible to *Bgt* #70 (Table S12).

The functional molecular marker *uhw403* was used for a large-scale screening of the distribution of *Pm69* among 538 wheat accessions of which 310 are WEW from across the Fertile Crescent and 228 represent other wheat species (Figure 5a, Table S13). Only G305-3M yielded positive PCR amplification. To further estimate the presence of the gene in the WEW gene pool, we went back to the original G305-3M collection site south of Kadita, Northern Israel, and collected additional 64 WEW accessions in a radius of less than 1km from the original collection site (Figure 5b and S14). Even there, we were able to find only three WEW accessions that gave amplification of *uhw403* marker and showed high resistance to *Bgt* #70 (Figure 5b-d, Table S14). Sanger sequencing confirmed that these accessions contain a functional *Pm69* allele identical to that of G305-3M. Moreover, eight resistant accessions were identified to contain *TdPm60* by marker *M-Pm60-S1*^37,43^, and additional five accessions contain unknown *Pm* genes. Therefore, we can conclude that *Pm69* is a rare NLR allele among natural WEW populations, even within its population of origin near the collection site of its donor accession G305-3M.

**Figure 5.**
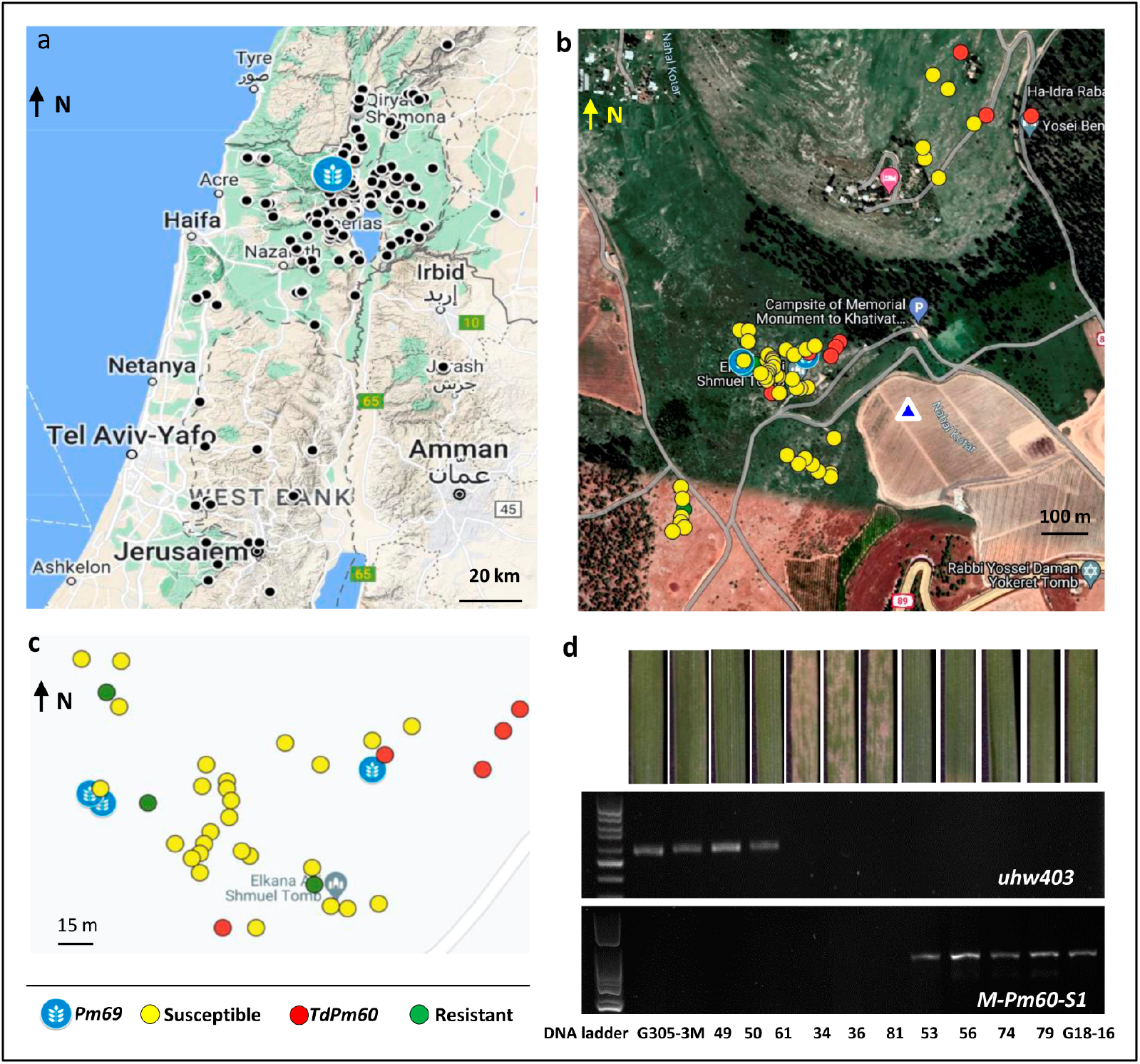
The geographic distribution of the *Pm69* allele in WEW populations and around the original G305-3M collection site south of Kadita, Northern Israel. **a** The geographic distribution of WEW natural populations (black circles) which were screened for the presence of *Pm69* functional allele. **b** The geographic distribution of WEW individual plants collected near the recorded G305-3M collection site (marked by a blue triangle). **c** A submap of b. The maps were obtained from Google Maps. **d** The Phenotypes in response to infection with *Bgt* #70 and agarose gel electrophoresis of PCR products amplified by marker *uhw403* (549 bp for *Pm69*) and *M-Pm60-S1* (831 bp for *TdPm60*) from representative WEW accessions (the ID information showed in Table S14).

### *Pm69* is located within a rapidly evolving NLR cluster

Micro-collinearity analysis among different wheat genomes revealed high copy number variation of NLRs in *Pm69* genetic regions, as well as multiple putative structural rearrangements (Figure 6 and S15). The WEW G305-3M harbors 47 NLRs and Zavitan 42, while all other domesticated wheat reference assemblies contain only 8-24 NLRs, within 0.5 - 3.1 Mb physical intervals comprising this cluster (Table S15). A highly similar *Pm69* homolog (NLR9, 92% similarity) was identified on contig utg5064 in this NLR cluster when searching for *Pm69* homologs in the ONT genome assembly of G305-3M (Table S12), suggesting a duplication event. Sequence alignment showed that NLRs in this cluster exhibited high diversity in the LRR domains relative to Pm69, while the Rx_N and part of NB-ARC domains were highly similar to Pm69 (Figure S16). Evolutionary analysis of all the functional R-proteins cloned in the *Triticeae* showed clustering of Pm69 with the leaf rust resistance protein Sr13 alleles (Similarity= 49%, Figure S17), originating from chromosome 6A of *T. turgidum* subsp. *dicoccum*^44,45^. Collinearity analysis indicates that *Pm69* and *Sr13* belong to homoeologous NLR clusters residing within syntenic regions located in the distal parts of group 6 chromosomes (Figure 6).

**Figure 6.**
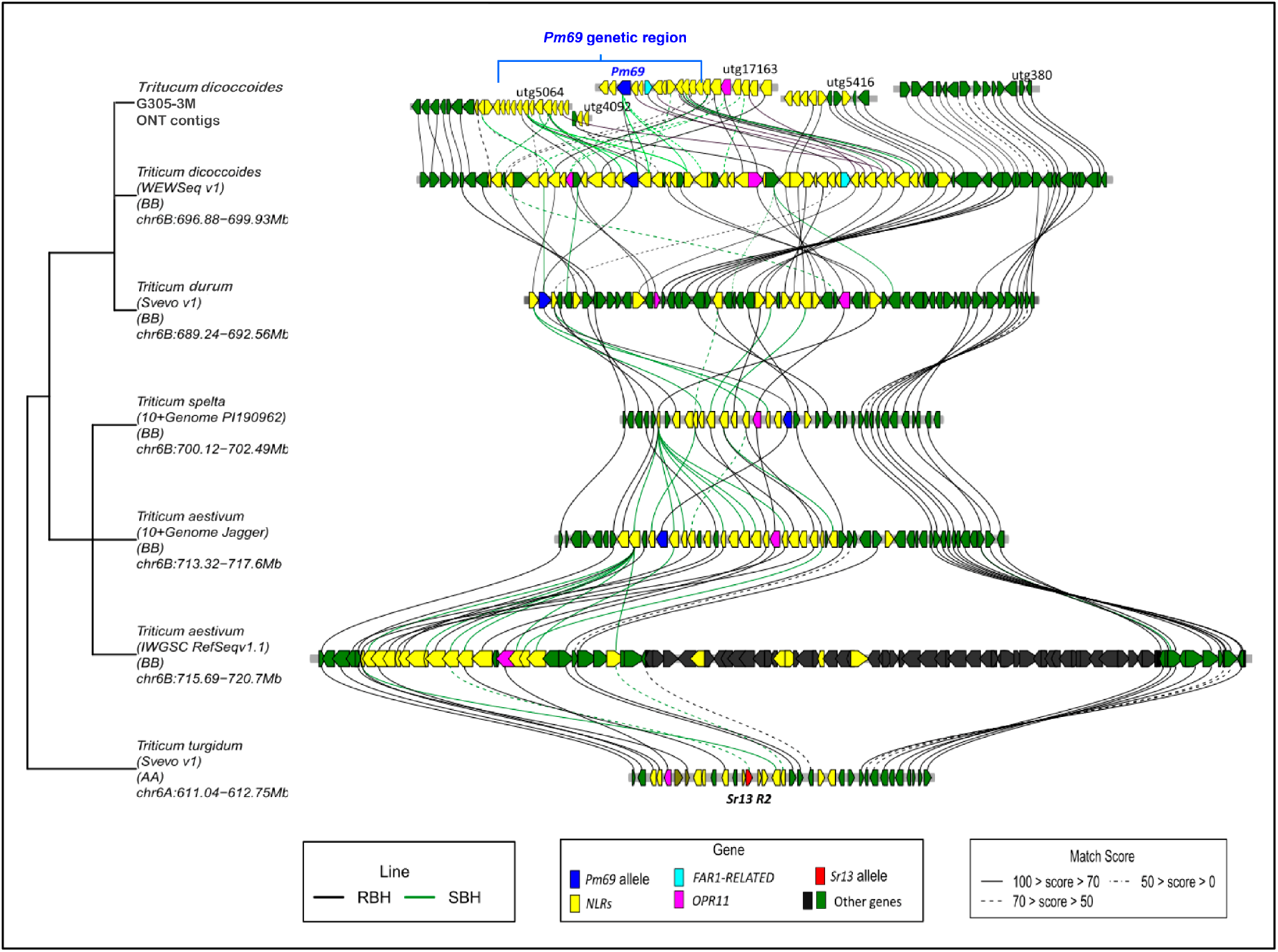
Micro-Collinearity analysis of the *Pm69* genetic region among different *Triticeae* genomes. The gene order of WEWseq_v1.0 was reversed according to the new version WEWseq_v2.0. Lines indicate similarity among genes. The blue genes represent the locations of *Pm69* homologs, the pink color is marking *OPR11* homologs, and the red genes represent *Sr13* R2 in Svevo A sub-genome (TRITD6Av1G225270, KY825226.1). RBH: Reciprocal best hit; SBH single-side best hit.

Inside the NLR cluster, 12-oxophytodienoate reductase 11 (OPR11), known to be involved in the biosynthesis of jasmonic acid (JA), showed a highly conserved protein sequence with 1-2 copies among different *Triticeae* genomes (Table S16), suggesting that opposing evolutionary forces are acting in a small genetic interval. Moreover, more distant regions flanking the clusters showed a high collinearity level, with similar content and order of genes. This might also imply that different evolutionary pressures may act on NLRs relative to other genes, probably imposed by natural selection (Figure 6).

### Introgression of *Pm69* gene into cultivated wheat and pyramiding with a yellow rust resistance gene

As part of the aim to develop resistant pre-breeding genetic resources, we transferred *Pm69* into elite Israeli bread wheat cultivar ‘Ruta’ using marker-assisted selection (MAS) following the “durum as a bridge” approach^36^ (Table S17, Figure 7). We also introgressed *Pm69* into the durum wheat cultivar ‘Svevo’, which contains *Sr13b*^44^. These homozygous ILs with different segments of G305-3M chromosome have been selected from the BC4F2 populations and showed high resistance to *Bgt* #70 (Figure 7a and Table S17).

**Figure 7.**
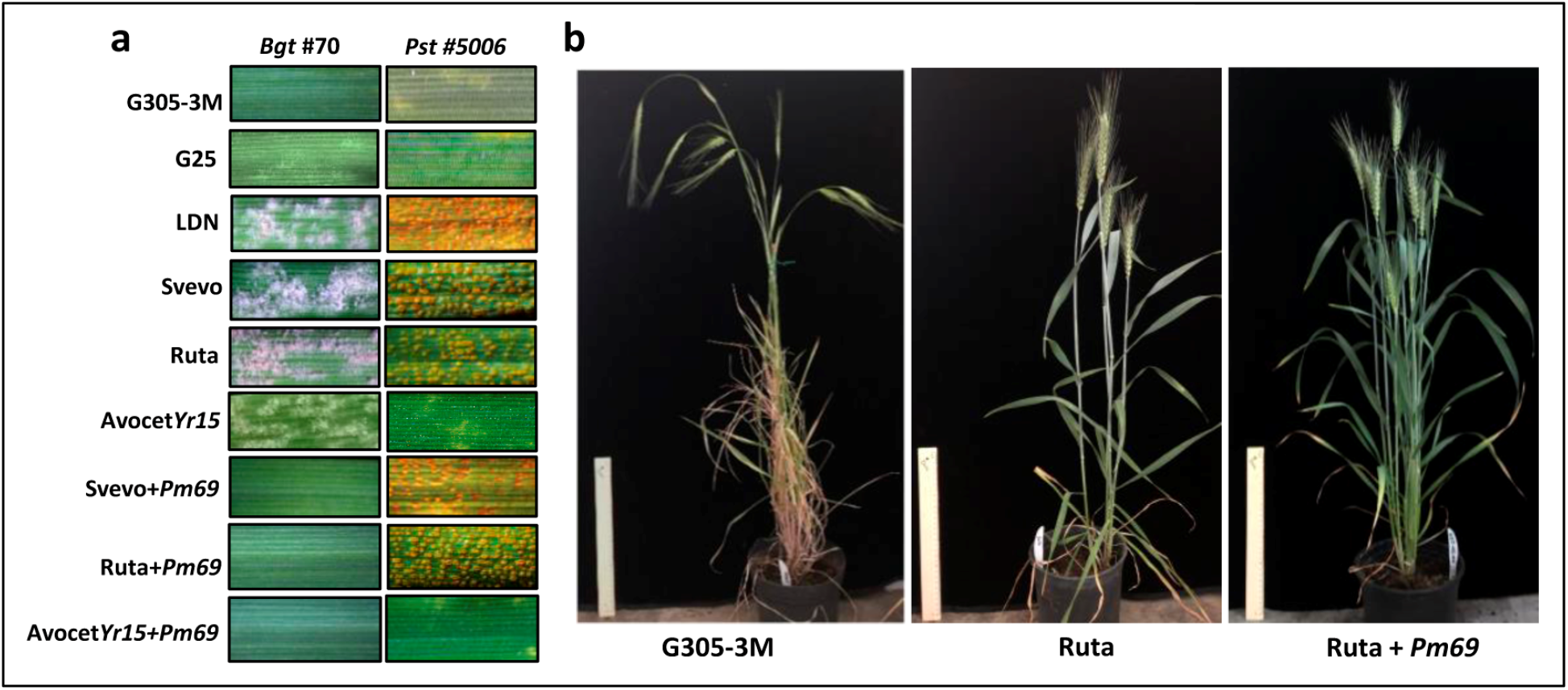
Introgression of the *Pm69* into the cultivated wheat and pyramiding with yellow rust resistance gene. **a** The phenotypes of different wheat parental and introgression lines to *Bgt* #70 and *Pst* #5006. WEW: G305-3M contains *Pm69* and G25 contains *Yr15*. Durum wheat: LDN and Svevo. Common wheat: Ruta and *AvocetYr15*. **b** The whole plant structure of the G305-3M, Ruta and a representative introgression line Ruta + *Pm69*.

Moreover, we found that G305-3M contains the yellow rust resistance gene *Yr15*, which was confirmed by our ONT sequencing data, present in 4.57 Mbp contig utg1161 and showed high resistance to *Pst* #5006 (Figure 7a). We pyramided *Pm69* with *Yr15* in the hexaploid cultivar ‘Avocet’ background, and selected the homozygous plants (Avocet*Yr15*+*Pm69*) by MAS, that showed high resistance to both *Pst* #5006 (IT = 1-3) and *Bgt* #70 (IT = 0;-1) (Figure 7a, Table S18). These resistant ILs could be employed in wheat resistance breeding programs in the future (Figure 7b).

## Discussion

The repertoire and diversity of *R* genes among crops’ wild relatives is a good source for resistance breeding against various diseases. The main strategies for protecting the wild relative’s gene pool are: (i) *in situ* conservation in nature (e.g. natural habitats) which allows them to continuously evolve novel resistances against rapidly evolving pathogens^46,47^, and (ii) *ex situ* conservation via on-site collections and longterm storage at gene banks (e.g. the Institute of Evolution Wild Cereal Gene Bank^48^). These collections can be used for identifying valuable genes for wheat breeding. To date, several genes have been already cloned from WEW *ex situ* collections, including the powdery mildew resistance genes *Pm41* and *TdPm60* that encode NLR immune receptors^8,9,37^, the yellow rust resistance genes *Yr15 (WTK1*) that encodes a tandem kinase-pseudokinase protein domains and *Yr36 (WKS1*) that encodes a protein with kinase and START lipid-binding like domains^49,50^. In the current study, we cloned the *Pm69* gene from a WEW accession G305-3M which confers wide-spectrum resistance to a worldwide collection of 55 *Bgt* isolates (Table S1). The *Pm69* is a rare NLR allele, located within a rapidly evolving NLR cluster, found in only one out of hundreds of WEW accessions available in our gene bank. This discovery of this new resistance gene is a highly valuable source for wheat resistance breeding.

The cloning of *Pm69* encountered several obstacles that are commonly faced by researchers attempting to pursue *R* gene cloning in wheat. The massive structural rearrangements in the *Pm69* genetic region suppressed the recombination rate and hampered the map-based cloning approach (Figure 2 and S2). An alternative approach we tried was the MutChromSeq for isolating and sequencing complete 6B chromosomes from G305-3M, the tetraploid *Pm69* donor. However, we had to abandon this attempt due to the low purity of the 6B chromosomes in fractions flow-sorted from the tetraploid wheat background (Figure S5). Eventually, we overcame the complexity of this rearranged locus by employing the ONT long-read sequencing technology to sequence the whole genome of the *Pm69* donor line G305-3M. Moreover, the MutRNA-seq approach helped us to identify a *Pm69* candidate within an NLR cluster by sequencing the transcriptome of the resistant wild-type G305-3M and its susceptible derivative mutants (Figure 3). MutRNA-seq approach is a useful method for identifying candidate genes based on comparisons of a wild-type line with its derivative mutants^51–53^. The main advantage of this combination of long-read sequencing and RNAseq method is that it allows to anchor short DNA reads to the long ONT contigs allowing to determine if two fragments belong to the same gene or separate genes. Furthermore, this method is suitable and solid for cloning of any type of gene for which mutants with clear phenotype can be generated, not necessarily R-genes (e.g. ONT-MutRNAseq, Figure S19).

The assembly of repetitive DNA and duplicated gene clusters, such as NLRs, is highly challenging when relying on short-read sequencing technologies^54^. The *Pm69* genetic region in IWGSC RefSeq v1.0 showed a large inversion compared with other reference genomes (Figure 6). However, this inversion was shown to be a result of an erroneous assembly, which has been corrected in the second version of this assembly (IWGSC RefSeq v2.1) obtained by PacBio long-read sequencing^27,55^ (Table S15). Similarly, the chromosome-scale assembly of a bread wheat cv. Kariega obtained by PacBio sequencing (34-fold genome) contributed to the cloning of *Yr27*^33^. Previously, the ONT sequencing was used to construct the *Crr3* locus in *Brassica napus* cv. Tosca and identify a duplicated TIR-NBS-LRR candidate for clubroot disease resistance gene^56^. In the current study, ONT long reads helped to resolve the complexity of the structural rearrangements within the rapidly evolving NLR cluster in the *Pm69* gene region. Therefore, with the drop of sequencing cost, long-read sequencing technologies are becoming the preferred method of choice in gene mapping and cloning, pan-genome era, and improving the assemblies of complex reference genomes, such as wheat^30^.

Gene duplications that create gene clusters are common characteristics of rapidly evolving genes, such as NLRs, often appearing to be the products of tandem duplication events, sometimes following unequal crossing over, as well as intra-cluster chromosomal rearrangements and gene conversion events^57^. For example, the exceptionally large size of *Phytophthora* resistance gene *Rps11* (27.7 kb) of soybean resulted from several rounds of inter- and intra-specific unequal recombination, located in a genomic region harboring a cluster of large NLR genes (7-12 NLRs)^58^. In wheat, powdery mildew resistance gene *MlWE74* on chromosome arm 2BS derived from WEW was also identified in an NBS-LRR gene cluster with 3-5 NLRs among different reference wheat genomes^59^. Here, we found more than 40 NLRs present in the *Pm69* genetic region (1.93-2.7M bp) of chromosome 6B in the WEW accessions (G305-3M and Zavitan), which are about two-fold higher numbers of NLRs as compared to domesticated wheat relatives (Table S15). The diversification of NLR gene numbers in different wheat species reflects the rapid evolutionary dynamics of plant R-genes along a relatively short evolutionary time^60^.

NLR gene cluster is found not only in the *Pm69* locus on wheat chromosome arm 6BL but also in the colinear region on chromosome arm 6AL, which contains the stem rust resistance gene *Sr13*^45^ within a cluster of 12 NLRs (Figure 6). These findings may indicate that these NLR clusters originated from the same diploid ancestor and continue to evolve since their split into A and B genome ancestors, resulting in at least two functional genes that confer resistance to different pathogens. Orthologous resistance genes from different *Triticeae* species can take different routes towards functionally active genes, as demonstrated by the two rye genes, *Pm17* and *Pm8*, located on chromosome arm 1RS that are orthologous or close paralogs of wheat *Pm3* located on wheat chromosome arm 1AS^2,10,61^. These different alleles might contribute to resistance to different pathogen isolates during the rapid co-evolution of *R-Avr* gene pairs^62^. The *Pm69* NLR cluster showed high polymorphism, while the *OPR11* located inside the cluster, as well as the further flanking regions of the cluster, were much more conserved among different wheat genomes. This might suggest that opposite evolutionary selection forces are acting on this genetic interval (Figure 6).

*Yr15* and *Pm69* genes, originated from two WEW accessions, G-25 and G305-3M, both collected by the late Dr. Gerechter-Amitai, in the upper Galilee, Israel, less than 5km apart. In a retrospective of four decades, this *ex-situ* collection and disease resistance identification enabled and led to the isolation and cloning of *Yr15* and *Pm69*^50,63^. The cloning of these genes not only provides solid evidence for the potential contribution of wild species to global food security but also prioritizes the *ex-situ* conservation of this exotic germplasm in gene banks, since natural habitats of WEW are becoming spatially fragmented and decreased due to the intense development and growth of human infrastructure. We were very lucky to still find two WEW plants with *Pm69* in the natural field, although the recorded G305-3M collection natural land site was destroyed by human structures (Figure 5b). However, human structures continue to invade this land and we may soon lose the *Pm69* resources forever without ex-situ conservation.

Long-term conservation of this genetic reservoir by the scientific community will assure further exploitation of useful new target genes and their introduction into cultivated crops via breeding. However, many genes that were introgressed into cultivated crops from their wild ancestors could not be easily used in plant breeding due to negative linkage drag and possible collateral damage associated with NLR diversity. In some cases, NLR genes can backfire and lead to genetic incompatibilities, often resulting in hybrid necrosis such as *Ne2* (*Lr13*)^64,65^. In wheat, specific combinations of Pm alleles (NLRs) can suppress each other, such as in the case of rye-derived *Pm8* which is suppressed by its wheat ortholog *Pm3*^66^. In the current study, we introgressed *Pm69* into the susceptible bread wheat cultivar ‘Ruta’ and pyramided it with the stripe rust R-gene *Yr15*. We also transferred *Pm69* into *T. durum* cv. Svevo background that contains a functional *Sr13* (Figure 7a). However, these *Pm69* introgression lines (ILs) combined with the functional NLRs or WTK genes, showed high resistance to *Bgt*, or *Bgt* and *Pst* without any obvious evidence of suppression (Figure 7a, Table S18). Considering the wide spectrum of resistance to *Bgt* isolates (Table S1), *Pm69* has great potential for future wheat resistance breeding.

## Methods

### Plant materials

Wild emmer wheat accession G305-3M (Accession number: CGN19852, https://www.genesys-pgr.org/10.18730/1NA3M), the *Pm69 (PmG3M*) gene donor line, was collected from Upper Galilee, Israel. G305-3M was crossed with the susceptible *T. durum* wheat line Langdon (LDN) to generate segregating mapping populations. RIL population was used for the construction of a sub-centiMorgan (cM) genetic linkage map. Differential wheat lines carrying different *Pm* genes (*Pm21, TdPm60* and *Pm30*) were used for testing the virulence of *Bgt* isolate #70. A core collection of WEW accessions, originating from the Fertile Crescent and obtained from the Wild Cereals Gene Bank (ICGB) at the University of Haifa, Israel, was used to screen the *Pm69* alleles. Modern Israeli bread wheat cultivar ‘Ruta’ and durum wheat cultivar ‘Svevo’, highly susceptible to *Bgt #70*, were used as recurrent parents for the development of hexaploid and tetraploid introgression lines. Hexaploid wheat lines Avocet+Yr15 that harbors *Yr15* were used as *Yr* gene donor lines or recurrent parents for pyramiding *Pm69* with *Yr* genes^50,67^.

### *Bgt* and *Pst* inoculation and disease assessment

*Bgt* #70 showed high virulence on many cultivated *T. durum, T. aestivum* and *T. monococcum* accessions (Figure S3), as well as on several *Pm* genes (e.g., *Pm1a-b, Pm2, Pm3a-d, Pm5a-b, Pm6, Pm7, Pm17, Pm22, Pm30* and *Pm1* + *Pm2* + *Pm9*)^68^, but it is avirulent on *Pm69*. Therefore, *Bgt* #70 was used in the current study for phenotyping of the *Pm69* mapping populations and EMS mutants. The *Bgt* inoculation process and the infection types (IT) were described in previous study^37^. The reactions to *Bgt* inoculation were examined visually, with the infection type (IT) recorded on a 0-4 scale at 7 days post-inoculation (dpi). Plants with an IT 0-2 (no disease colonies or less than 1 mm) were considered resistant and those with IT 3-4 (scattered disease spots of greater than 1 mm) are susceptible. Israeli *Pst* isolate #5006 was used for the yellow rust resistance test as described in previous study^50^. It was scored at 14 dpi using a 0-9 scale: 0-3 = resistant, none to trace sporulation; 4-6 = intermediate, light to moderate sporulation; 7-9 = susceptible, abundant sporulation. ROS accumulation and cell death were evaluated in inoculated wheat leaves (3 dpi) as described by our previous study^37^.

### Development of molecular markers, genetic and physical maps

Plant genomic DNA was extracted using the CTAB method from leaves of 2-3 weeks-old wheat seedlings. The primary genetic maps of *Pm69* using cleaved amplified polymorphic sequence (CAPS) and Sequence-Tagged Site (STS) markers were developed based on the local synteny of wheat with major cereals like rice, *Brachypodium, Sorghum* and barley genomes^69^. For high-resolution mapping we used Kompetitive Allele Specific PCR (KASP) markers developed using PolyMarker (http://polymarker.tgac.ac.uk/) based on specific SNPs identified by the wheat 90K iSelect SNP array and the wheat 15K SNP array (Trait Genetics®, Germany). The International Wheat Genome Sequencing Consortium (IWGSC) RefSeq assembly v1.0^27^, Wild Emmer Genome Assembly (Zavitan WEW_v.1.0, WEW_v2.0)^40,70^ and the Durum Wheat Svevo RefSeq Rel. 1.0^41^ were used to design SSR markers by the BatchPrimer3 website tools, as well as for development of KASP markers after verifying polymorphism between the parental lines by Sanger sequencing of PCR products. PCR reactions were performed by using 2×Taq PCR Master Mix (TIANGEN, China). KASP marker analysis was performed on the StepOnePlus Real-Time PCR system (Applied Biosystems, USA) as described by a previous study^37^.

*Pm69*-flanking markers (*uhwk386* and *uhwk399*) were used to select 147 RILs that carry critical recombination events in the *Pm69* gene region after screening of 5500 F2 plants (G305-3M ×LDN). These RILs were used for further dissection of the target locus by using the graphical genotyping approach^71^. The genetic distances between the detected markers and *Pm69* were calculated based on genotypic and phenotypic data using JoinMap 5.0. The physical location of the primers of each marker was identified by BLAST + search against the sequences of the chromosome arm 6BL of wheat reference genomes. Genetic and physical maps of *Pm69* were constructed using MapChart v2.2.

### Mutant development

To produce loss-of-function mutants of the *Pm69* gene, we mutagenized the wild emmer accession G305-3M and homozygous resistant F6 RIL-(169B) by using 0.3%-0.5% Ethyl methane sulfonate (EMS) treatments, as described by previous study^72^. We generated 1530 G305-3M and 162 169B M0 plants which were developed to M2 families. We inoculated 10-20 plants of each M2 family with *Bgt* #70 for screening for susceptible mutants under controlled greenhouse conditions. Susceptible mutants and their resistant sister lines were selected and grown to M3-M4 generation. Resistant sister lines were kept as controls for verification of the *Pm69* candidate gene. Susceptible M3-M4 independent mutants, one from RIL 169B and four from G305-3M EMS-treated populations, were selected and confirmed with 30 *Pm69* flanking markers that showed the same haplotype as G305-3M, ruling out the possibility of cross-pollination from other susceptible lines.

### Chromosome sorting

Mitotic metaphase chromosome suspensions were prepared from tetraploid wheat lines G305-3M and LDN, and from hexaploid introgression wheat line SC28RRR-26 following by previous research^73,74^. Briefly, cell cycle of meristematic root tip cells was synchronized using hydroxyurea, and mitotic cells were accumulated in metaphase using amiprohos-methyl. Suspensions of intact chromosomes were prepared by mechanical homogenization of 100 formaldehyde-fixed root tips in 600 μl LB01 buffer^75^. GAA microsatellites and/or GAA and ACG microsatellites were labelled on chromosomes by fluorescence *in situ* hybridization in suspension (FISHIS) using FITC-labelled oligonucleotides (Sigma, Saint Louis, USA) as has been shown before^76^ and chromosomal DNA was stained by DAPI (4’,6-diamidino 2-phenylindole) at 2 μg/ml. Bivariate flow karyotypes FITC vs. DAPI fluorescence were acquired using a FACSAria II SORP flow cytometer and sorter (Becton Dickinson Immunocytometry Systems, USA). The samples were analyzed at rates of 1,500–2,000 particles and different positions of sorting windows were tested on bivariate flow karyotype FITC vs. DAPI to achieve the highest purity in the sorted 6B fractions. The content of flow-sorted fractions was estimated using microscopy analysis of slides, containing 1,500–2,000 chromosomes, sorted into a 10-μl drop of PRINS buffer^77^. Sorted chromosomes were identified by FISH with probes for DNA repeats pSc119.2, Afa family and 45S rDNA according to previous study^78^. At least 100 chromosomes were classified for each sample using a standard karyotype^73^.

### Nanopore and Illumina sequencing

*DNA extraction and QC:* High molecular weight (HMW) DNA was extracted from isolated nuclei and purified following a modified salting out DNA extraction protocol (10X Genomics)^79^. Stock HMW DNA was size selected on a Blue Pippin instrument (Sage Science) with the high pass protocol and electrophoretic conditions to retain fragments > 30 kb. Eluate was bead cleaned and concentrated. Size selected DNA’s were quantified by fluorometry (Qubit 2.0) and DNA integrity was evaluated using a Tapestation 2200 instrument (Agilent). HMW DNA’s were stored at 4°C until library preparation. *Library preparation:* Long molecule libraries (1D: Oxford Nanopore Technologies) were prepared following the standard 1D ligation protocol (LSK109) with minor modifications to retain and enrich for HMW molecules. Briefly 1.2μg of size selected, end-repaired HMW DNA’s were used as input into each library preparation reaction. Libraries were sequenced with R9 flow cell on a PromethION instrument with high accuracy base-calling enabled. Raw read data were filtered for size and quality score, and approximately 305 Gb filtered data (~23X coverage) was used for assembly. Illumina DNA prep libraries were prepared, indexed, and sequenced to ~25X coverage on the NovaSeq 6000 S4 flow cell for polishing (PE reads: 150 bp).

### Genome assembly, polishing, and scaffolding

Raw fast5 files generated by ONT sequencing of G305-3M were base-called using Guppy version 3.6 (Oxford Nanopore Technology) to produce fastq files. Fastq files from multiple flow-cells were concatenated to form a consolidated fastq file, which was then used for genome assembly using Smartdenovo with the default parameters^80^. The raw assembly from Smartdenovo was then subjected to a single round of long read polishing using Medaka version 144 (https://github.com/nanoporetech/medaka) followed by two rounds of short read polishing using Pilon (Walker et al., 2014). Assembly stats were calculated using QUAST version 5.0.2 (https://academic.oup.com/bioinformatics/article/29/8/1072/228832)

### Construction of *Pm69* physical map and comparison with other reference genomes

The co-dominant markers in the *Pm69* genetic region and candidate genes from WEW_v2.0 were used for searching the contigs of G305-3M ONT assembly. The contigs that showed a perfect match with these markers were selected for the construction of the *Pm69* physical map. The new mapping markers were developed based on the polymorphism between the ONT contigs and wheat reference genomes. Those markers were used to genotype the 31 representative RILs to characterize the size of the chromosome region that is co-segregating with *Pm69*. The G305 ONT contigs were sliced into 1Mb segments with pyfasta and mapped to the wild emmer assembly WEW_v2.0 using BWA-MEM software^81^. The G305-3M ONT contigs anchored to the *Pm69* region were compared to the Zavitan (WEW_v2.0) and Svevo RefSeq Rel. 1.0 genomes using Minimap (https://github.com/lh3/minimap2), and visualized using ggplot geom_segment in R platform.

### Transcriptome sequencing

Wheat leaf samples from the four susceptible mutants and the G305-3M resistant wild type were collected 24 hours after 0.1 mM 2,1,3-Benzothiadiazole (BTH) treatment with 0.05% Tween-20. Treated leaves were preserved in RNAlater (Sigma-Aldrich, UK) and sent for RNA extraction and sequencing in Novogene-UK. Samples were sequenced on Illumina NovaSeq 6000 instrument. About 40 million 150 bp paired-end (PE) reads were obtained for each sample. Those RNAseq reads were aligned to the G305-3M ONT genomic DNA contigs using GSNAP with default settings followed by sorting^82^, duplicate removal, and indexing using SAMtools. Mutation detection was done by visualizing the obtained bam files with IGV genome browser focusing on the contigs that were anchored to the *Pm69* region^83^. We looked for the regions on the contigs that had >4 read depth along at least 1kb and had mutations in all susceptible mutant samples relative to the resistant G305-3M wild-type RNA reads and to the reference G305-3M genome. The mutations had to be in all reads covering the site and to have read quality >30. Moreover, *de novo* assembly of the wild-type RNAseq reads were obtained using TRANSABYSS software^84^.

### Gene annotations, cloning of *Pm69* and Sanger sequencing

Gene annotation of the ONT contigs mapped to *Pm69* flanking region was carried on GeneSAS 6.0 server (https://www.gensas.org/gensas) using BLAST for EST matching and Augustus for structural prediction. Annotation of resistance genes was done using NLR annotator^85^, PfamScan (ftp://ftp.ebi.ac.uk/pub/databases/Pfam/Tools/), Fgenesh gene-finder (http://www.softberry.com/berry.phtml) and NCBI BLASTP. The primers for cloning and sequencing of *Pm69* were designed based on ONT contig utg17163 sequences and the RNA-seq assembly using BatchPrimer3 website tools. *Pm69* was amplified from cDNA library of G305-3M using the VeriFi™ Polymerase (PCRBIO, UK), then inserted into a cloning plasmid using 5 min™ TA/Blunt-Zero Cloning Kit (Vazyme, China), and transformed into *E. coli* strain DH5α. The plasmid was extracted by Hybrid-Q™ Plasmid Rapidprep Kit (GeneAll, Korea) and sequenced using BigDye Terminator v1.1 Cycle Sequencing Kit (Applied Biosystems, USA) on ABI 3130 instrument (Applied Biosystems, USA).

### RNA extraction and quantitative Real-Time-PCR (qRT-PCR)

Total RNA was extracted from *Bgt* #70-inoculated, and non-inoculated G305-3M leaf segments collected along 10 different time points (0, 3, 6, 9, 12, 16, 24, 36, 48, 72 hpi) using the RNeasy Plant Mini Kit (Qiagen, Germany). The cDNA was synthesized from total RNA using a qScript™ cDNA Synthesis Kit (Quantabio, USA). The gene-specific primers of the *Pm69* and the housekeeping gene *Ubiquitin* were used for qRT-PCR amplification performed on a StepOne thermal cycler (ABI, USA) in a volume of 10 μl containing 5 μl of SYBR Green FastMix (Quantabio, USA), 250 nM primers and optimized dilution of cDNA template. The program included an initial step at 95 °C for 30 s followed by 40 cycles of 95 °C for 15 s, 60 °C for 30 s, and 72 °C for 10 s. Relative expression of the target genes was calculated by 2^^(ubiquitin CT-Target CT)^ ± standard error of the mean (SEM). All of the qRT-PCRs were performed in triplicate, each with at least three (up to five) independent biological repetitions.

### Virus-induced gene silencing (VIGS)

The construction of VIGS vectors and inoculation were carried out as described earlier^86^. To specifically silence the *Pm69* gene without off-targets in the G305-3M transcriptome, we used si-Fi software to predict 150-350 nt gene regions with the efficient siRNAs^87^. Two selected regions were amplified and inserted in the *pCa-BSMV-γ* vector, as well as the control insertions of *GFP* and *PDS* genes. Equimolar amount of *Agrobacterium tumefaciens* strain GV3101 with *pCa-BSMV-α, pCa-BSMV-β* and *pCa-BSMV-γ* vectors carrying target or control genes were used to inoculate *Nicotiana benthamiana* leaves to produce virus transcripts. Infiltrated *N. benthamiana* leaves were then used to extract the sap and further inoculate the second leaves of two-weeks old wheat plantlets. When the pCa-BSMV-γ-PDS silenced plants showed chlorosis in the leaves, those plants were inoculated with *Bgt* #70 in a growth chamber. Two weeks post inoculation, the reaction of wheat plants to *Bgt* inoculation was recorded.

### Phylogeny and Synteny analysis

The evolutionary tree was built using MEGA-X (aligning by MUSCLE, Neighbor-Joining algorithm, bootstrap: 500 times) and the tree was draw using iTOL v6 (https://itol.embl.de/). Synteny was analyzed by TGT http://wheat.cau.edu.cn/TGT/. Pm69 homologs were identified by blasting the Pm69 protein sequence to the database of WheatOmics 1.0, EnsemblPlants, and NCBI. The alignment of Pm69 homologs was done by the multiple alignment tool CLUSTALW from NCBI and displayed using ESPript (https://espript.ibcp.fr/ESPript/cgi-bin/ESPript.cgi).

### Statistical analysis

Statistical analysis was performed using JMP® version 15.1 statistical packages (SAS Institute, Cary, NC, USA). Comparison between the treatments were performed by Student’s t-test. Asterisks indicate the level of significance at p < 0.05 (*), p < 0.01 (**) and p < 0.001 (***).

## Data availability

The ONT sequencing contigs and raw reads of the MutRNAseq were deposited to the NCBI Sequence Read Archive under BioProject ID PRJNA795708. Correspondence and requests for materials and data should be addressed to Tzion Fahima.

## Acknowledgments

The authors wish to thank Dr. Olga Borzov, Dr. Tamar Kis-papo, Dr. Tamar Krugman and Ms. Souad Khalifa for their professional and moral support, Dr. Mahmoud Said, Zdeňka Dubská and Jitka Weiserová for their assistance with chromosome sorting, Dr. Imad Shams and Dr. Tamar Lotan Labs for assistance in microscopy work, Dr. Amir Raz and Prof. Avi Levy for providing the VIGS plasmids and *Nicotiana benthamiana* seeds, Andrew G Sharpe and Chu Shin Koh for their assistance with raw ONT assembly, and Dr. Lin Huang for assistance in screening for the distribution of *Pm69* in wheat accessions. This study was supported by the Israel Science Foundation, grant numbers 1366/18 and 2342/18 and the Genome Canada funded project 4D Wheat (CJP). IM and JD were supported by ERDF project Plants as a Tool for Sustainable Global Development (No.CZ.02.1.01/0.0/0.0/16_019/0000827). Dr. YL was supported by a fellowship provided by the Planning and Budgeting Committee (PBC) of the Israel Council for Higher Education for Outstanding Post-doctoral Fellows from China and India.

## Author contributions

TF, CP, HS, and YL conceived and designed the study; ZW, YL, RBD, VB and VK developed the critical RILs and genetic mapping; ZW, YL and SJ developed and screened the mutants. JD and IM performed chromosome flow sorting. CJP, HSC, JE and KW performed ONT sequencing and assembly. YL, HS and RR performed the MutRNA-seq. YL and RR performed qPCR and *Pm69* cloning. YL, HS and LG did the VIGS. YL and ZW developed the introgression lines. YL and HS conducted the evolutionary analysis; YL, ZW, SJ, LG, and VK performed the phenotyping of *Bgt* and *Pst* resistance. LY, PBP, ZW, and HS analyzed *Pm69* alleles in the different wheat accessions. YL, TF, HS, and ZW drafted the manuscript; TF, CP, JD, LG, VK, HSC, JE, KW, RBD, and RR reviewed and edited the manuscript; TF was responsible for coordination and funding acquisition.

## Additional information

### Extended data

is available for this paper at XXXXXXXXXXXXXXXXXXXXXX.

### Supplementary information

The online version contains supplementary material available at XXXXXXXXXXXXXXXXXXXXXXXX

### Correspondence and requests for materials and data

should be addressed to Tzion Fahima.

## References

1. McIntosh R. A., Dubcovsky J., Rogers W. J., Xia X. C. & Raupp W. J. Catalogue of gene symbols for wheat: 2019 supplement. (2019).

2. Hurni, S. et al. Rye *Pm8* and wheat *Pm3* are orthologous genes and show evolutionary conservation of resistance function against powdery mildew. Wiley Online Library 76, 957–969 (2013).

3. Sánchez-Martín, J. et al. Rapid gene isolation in barley and wheat by mutant chromosome sequencing. Genome Biol 17, (2016).

4. Xing, L. et al. Pm21 from Haynaldia villosa encodes a CC-NBS-LRR protein conferring powdery mildew resistance in wheat. Molecular Plant, 11(6), 874–878 (2018).

5. Zou, S. et al. The NB-LRR gene *Pm60* confers powdery mildew resistance in wheat. Wiley Online Library 218, 298–309 (2018).

6. Xie, J. et al. A rare single nucleotide variant in *Pm5e* confers powdery mildew resistance in common wheat. Wiley Online Library 228, 1011–1026 (2020).

7. Hewitt, T. et al. A highly differentiated region of wheat chromosome 7AL encodes a Pm1a immune receptor that recognizes its corresponding AvrPm1a effector from Blumeria. Wiley Online Library 229, 2812–2826 (2021).

8. Wu, Q. et al. Functional characterization of powdery mildew resistance gene *MlIW172*, a new *Pm60* allele and its allelic variation in wild emmer wheat. Journal of Genetics and Genomics 49, 787–795 (2022).

9. Li, M. et al. A CNL protein in wild emmer wheat confers powdery mildew resistance. New Phytologist, 228(3), 1027–1037 (2020).

10. Yahiaoui, N., Srichumpa, P., Dudler, R. & Keller, B. Genome analysis at different ploidy levels allows cloning of the powdery mildew resistance gene Pm3b from hexaploid wheat. The Plant Journal 37, 528–538 (2004).

11. Lu, P. et al. A rare gain of function mutation in a wheat tandem kinase confers resistance to powdery mildew. Nature Communications 2020 11:1 11, 1–11 (2020).

12. Sánchez-Martín, J. et al. Wheat *Pm4* resistance to powdery mildew is controlled by alternative splice variants encoding chimeric proteins. Nat Plants 7, 327–341 (2021).

13. Krattinger, S. G. et al. A putative ABC transporter confers durable resistance to multiple fungal pathogens in wheat. Science (1979) 323, 1360–1363 (2009).

14. Moore, J. et al. A recently evolved hexose transporter variant confers resistance to multiple pathogens in wheat. Nature genetics, 47(12), 1494–1498 (2015).

15. Golzar, H., Shankar, M. & D’Antuono, M. Responses of commercial wheat varieties and differential lines to western Australian powdery mildew *(Blumeria graminis* f. Sp. *tritici)* populations. Australasian Plant Pathology 45, 347–355 (2016).

16. Walkowiak, S. et al. Multiple wheat genomes reveal global variation in modern breeding. Nature, 588(7837), 277–283 (2020).

17. Bohra, A. et al. Reap the crop wild relatives for breeding future crops. Trends Biotechnol (2021).

18. Dong, L. et al. Rapid evolutionary dynamics in a 2.8-Mb chromosomal region containing multiple prolamin and resistance gene families in Aegilops tauschii. The Plant Journal 87, 495–506 (2016).

19. Andersen, E. J. et al. Wheat disease resistance genes and their diversification through integrated domain fusions. Front Genet 898 (2020).

20. Shizuya, H. et al. Cloning and stable maintenance of 300-kilobase-pair fragments of human DNA in Escherichia coli using an F-factor-based vector. National Acad Sciences 89, 8794–8797 (1992).

21. Zhang, J., Zhang, P., Dodds, P. & Lagudah, E. How Target-Sequence Enrichment and Sequencing (TEnSeq) Pipelines Have Catalyzed Resistance Gene Cloning in the Wheat-Rust Pathosystem. Front Plant Sci 11, (2020).

22. Steuernagel, B. et al. Rapid cloning of disease-resistance genes in plants using mutagenesis and sequence capture. Nature biotechnology, 34(6), 652–655 (2016).

23. Arora, S. et al. Resistance gene cloning from a wild crop relative by sequence capture and association genetics. Nat Biotechnol 37, 139–143 (2019).

24. Thind, A. et al. Rapid cloning of genes in hexaploid wheat using cultivar-specific long-range chromosome assembly. Nature Biotechnology, 35(8), 793–796 (2017).

25. Molnár, I. et al. Flow cytometric chromosome sorting from diploid progenitors of bread wheat, T. urartu, Ae. speltoides and Ae. tauschii. Theoretical and Applied Genetics 127, 1091–1104 (2014).

26. Wu, Q. et al. Bulked segregant CGT-Seq-facilitated map-based cloning of a powdery mildew resistance gene originating from wild emmer wheat (Triticum dicoccoides). Plant Biotechnol J 19, 1288 (2021).

27. (IWGSC), I. W. G. S. C. et al. Shifting the limits in wheat research and breeding using a fully annotated reference genome. Science (1979) 361, eaar7191 (2018).

28. Rhoads, A. et al. PacBio sequencing and its applications. Genomics, proteomics & bioinformatics, 13(5), 278–289 (2015).

29. Deamer, D., Akeson, M. & Branton, D. Three decades of nanopore sequencing. Nature Biotechnology 2016 34:5 34, 518–524 (2016).

30. Aury, J. M. et al. Long-read and chromosome-scale assembly of the hexaploid wheat genome achieves high resolution for research and breeding. Gigascience 11, 1–18 (2022).

31. Mascher, M. et al. Long-read sequence assembly: a technical evaluation in barley. The Plant Cell, 33(6), 1888–1906 (2021).

32. Sato, K. et al. Chromosome-scale genome assembly of the transformation-amenable common wheat cultivar ‘Fielder’. DNA Research, 28(3), dsab008 (2021).

33. Athiyannan, N. et al. Long-read genome sequencing of bread wheat facilitates disease resistance gene cloning. Nat Genet 1–5 (2022).

34. Amarasinghe, S. L. et al. Opportunities and challenges in long-read sequencing data analysis. Genome Biol 21, 1–16 (2020).

35. Nilsen, K. et al. Copy number variation of TdDof controls solid-stemmed architecture in wheat. National Acad Sciences 117, 28708–28718 (2020).

36. Klymiuk, V. et al. Durum wheat as a bridge between wild emmer wheat genetic resources and bread wheat. Applications of Genetic and Genomic Research in Cereals. Woodhead Publishing, 201–230 (2019).

37. Li, Y. et al. TdPm60 identified in wild emmer wheat is an ortholog of *Pm60* and constitutes a strong candidate for *PmG16* powdery mildew resistance. Theoretical and Applied Genetics 134, 2777–2793 (2021).

38. Xie, W. et al. Identification and characterization of a novel powdery mildew resistance gene *PmG3M* derived from wild emmer wheat, Triticum dicoccoides. Theoretical and Applied Genetics 124, 911–922 (2012).

39. Wei, Z. Z., Klymiuk, V., Bocharova, V., Pozniak, C. & Fahima, T. A post-haustorial defense mechanism is mediated by the powdery mildew resistance gene, *PmG3M*, derived from wild emmer wheat. Pathogens 9, (2020).

40. Avni, R. et al. Wild emmer genome architecture and diversity elucidate wheat evolution and domestication. Science 357, 93–97 (2017).

41. Maccaferri, M. et al. Durum wheat genome highlights past domestication signatures and future improvement targets. Nat Genet 51, 885–895 (2019).

42. Holzberg, S., Brosio, P., Gross, C. & Pogue, G. P. Barley stripe mosaic virus-induced gene silencing in a monocot plant. The Plant Journal 30, 315–327 (2002).

43. Zhao, F. et al. Powdery mildew disease resistance and marker-assisted screening at the Pm60 locus in wild diploid wheat Triticum urartu. Crop Journal 8, (2020).

44. Gill, B. et al. Function and evolution of allelic variations of *Sr13* conferring resistance to stem rust in tetraploid wheat *(Triticum turgidum* L.). The Plant Journal, 106(6), 1674–1691 (2021).

45. Zhang, W. et al. Identification and characterization of *Sr13*, a tetraploid wheat gene that confers resistance to the Ug99 stem rust race group. Proc Natl Acad Sci U S A 114, E9483–E9492 (2017).

46. Fu, Y. B. et al. Elevated mutation and selection in wild emmer wheat in response to 28 years of global warming. Proc Natl Acad Sci U S A 116, 20002–20008 (2019).

47. Huang, L. et al. Evolution and Adaptation of Wild Emmer Wheat Populations to Biotic and Abiotic Stresses. Annu Rev Phytopathol 54, 279–301 (2016).

48. Krugman, T., Nevo, E., Beharav, A., Sela, H. & Fahima, T. The Institute of Evolution Wild Cereal Gene Bank at the University of Haifa. Isr J Plant Sci 65, 129–146 (2018).

49. Fu, D. et al. A kinase-START gene confers temperature-dependent resistance to wheat stripe rust. Science (1979) 323, 1357–1360 (2009).

50. Klymiuk, V. et al. Cloning of the wheat *Yr15* resistance gene sheds light on the plant tandem kinase-pseudokinase family. Nature communications, 9(1), 1–12 (2018).

51. Yu, G. et al. Aegilops sharonensis genome-assisted identification of stem rust resistance gene Sr62. Nature communications, 13(1), 1–13 (2022).

52. Yu, G. et al. The wheat stem rust resistance gene Sr43 encodes an unusual protein kinase. doi:10.21203/rs.3.rs-1820134/v1 (2022).

53. Wang, Y. et al. An unusual tandem kinase fusion protein confers leaf rust resistance in wheat. doi:10.21203/RS.3.RS-1807889/V1 (2022).

54. Tørresen, O. K. et al. Tandem repeats lead to sequence assembly errors and impose multi-level challenges for genome and protein databases. Nucleic Acids Res 47, 10994–11006 (2019).

55. Zhu, T. et al. Optical maps refine the bread wheat Triticum aestivum cv. Chinese Spring genome assembly. The Plant Journal 107, 303–314 (2021).

56. Kopec, P. M. et al. Local Duplication of TIR-NBS-LRR Gene Marks Clubroot Resistance in Brassica napus cv. Tosca. Front Plant Sci 12, (2021).

57. Barragan, A. C. & Weigel, D. Plant NLR diversity: the known unknowns of pan-NLRomes. Plant Cell 33, 814–831 (2021).

58. Wang, W. et al. A giant NLR gene confers broad-spectrum resistance to *Phytophthora sojae* in soybean. Nature Communications 2021 12:1 12, 1–8 (2021).

59. Zhu, K. et al. Fine mapping of powdery mildew resistance gene *MlWE74* derived from wild emmer wheat (Triticum turgidum ssp. *dicoccoides)* in an NBS-LRR gene cluster. Theoretical and Applied Genetics 135, 1235–1245 (2022).

60. Bergelson, J., Kreitman, M., Stahl, E. A. & Tian, D. Evolutionary dynamics of plant R-genes. Science (1979) 292, 2281–2285 (2001).

61. Singh, S. P. et al. Evolutionary divergence of the rye *Pm17* and *Pm8* resistance genes reveals ancient diversity. Plant Mol Biol 98, 249–260 (2018).

62. Frantzeskakis, L. et al. Rapid evolution in plant–microbe interactions – a molecular genomics perspective. New Phytologist 225, 1134–1142 (2020).

63. Gerechter-Amitai, Z. K. & van Silfhout, C. H. Resistance to powdery mildew in wild emmer (Triticum dicoccoidesKörn.). Euphytica 33, 273–280 (1984).

64. Hewitt, T. et al. Wheat leaf rust resistance gene *Lr13* is a specific *Ne2* allele for hybrid necrosis. Molecular Plant, 14(7), 1025–1028 (2021).

65. Yan, X. et al. High-temperature wheat leaf rust resistance gene *Lr13* exhibits pleiotropic effects on hybrid necrosis. Molecular Plant, 14(7), 1029–1032 (2021).

66. Hurni, S. et al. The powdery mildew resistance gene *Pm8* derived from rye is suppressed by its wheat ortholog *Pm3*. The Plant Journal 79, 904–913 (2014).

67. Yaniv, E. et al. Evaluation of marker-assisted selection for the stripe rust resistance gene *Yr15*, introgressed from wild emmer wheat. Molecular Breeding 35, 1–12 (2015).

68. Ben-David, R. et al. Differentiation among *Blumeria graminis* f. sp. tritici isolates originating from wild versus domesticated *Triticum* species in Israel. Phytopathology 106, 861–870 (2016).

69. Raats, D. et al. Application of CAPS Markers for Genomic Studies in Wild Emmer Wheat. New York: Nova Science Publishers 31–60 (2014).

70. Zhu, T. et al. Improved Genome Sequence of Wild Emmer Wheat Zavitan with the Aid of Optical Maps. G3 Genes|Genomes|Genetics 9, 619–624 (2019).

71. Distelfeld, A., Uauy, C., Fahima, T., Phytologist, J. D.-N. & 2006, undefined. Physical map of the wheat high-grain protein content gene *Gpc-B1* and development of a high-throughput molecular marker. Wiley Online Library 169, 753–763 (2006).

72. Uauy, C. et al. A modified TILLING approach to detect induced mutations in tetraploid and hexaploid wheat. BMC Plant Biol 9, (2009).

73. Kubaláková, M. et al. Chromosome Sorting in Tetraploid Wheat and Its Potential for Genome Analysis. Genetics 170, 823–829 (2005).

74. Vrana, J. et al. Flow sorting of mitotic chromosomes in common wheat (Triticum aestivum L.). Genetics 156, 2033 (2000).

75. Dpooležel, J., Binarová, P. & Lcretti, S. Analysis of Nuclear DNA content in plant cells by Flow cytometry. Biologia Plantarum 1989 31:2 31, 113–120 (1989).

76. Giorgi, D. et al. FISHIS: Fluorescence In Situ Hybridization in Suspension and Chromosome Flow Sorting Made Easy. PLoS One 8, e57994 (2013).

77. Kubaláková, M., Macas, J. & Doležel, J. Mapping of repeated DNA sequences in plant chromosomes by PRINS and C-PRINS. Theoretical and Applied Genetics 1997 94:6 94, 758–763 (1997).

78. Molnár, I. et al. Dissecting the U, M, S and C genomes of wild relatives of bread wheat *(Aegilops* spp.) into chromosomes and exploring their synteny with wheat. The Plant Journal 88, 452–467 (2016).

79. Zhang, M. et al. Preparation of megabase-sized DNA from a variety of organisms using the nuclei method for advanced genomics research. Nat Protoc 7, 467–478 (2012).

80. Liu, H., Wu, S., Li, A. & Ruan, J. SMARTdenovo: A de novo Assembler Using Long Noisy Reads. doi:10.20944/PREPRINTS202009.0207.V1 (2020).

81. Li, H. Aligning sequence reads, clone sequences and assembly contigs with BWA-MEM. doi:10.48550/arxiv.1303.3997 (2013).

82. Wu, T. D., Reeder, J., Lawrence, M., Becker, G. & Brauer, M. J. GMAP and GSNAP for genomic sequence alignment: Enhancements to speed, accuracy, and functionality. Methods in Molecular Biology 1418, 283–334 (2016).

83. Robinson, P. & Zemo jtel, T. Integrative genomics viewer (IGV): Visualizing alignments and variants. Computational Exome and Genome Analysis 233–245 (2017).

84. Robertson, G. et al. De novo assembly and analysis of RNA-seq data. Nature Methods 2010 7:11 7, 909–912 (2010).

85. Steuernagel, B. et al. The NLR-Annotator Tool Enables Annotation of the Intracellular Immune Receptor Repertoire. Plant Physiol 183, 468–482 (2020).

86. Yuan, C. et al. A high throughput barley stripe mosaic virus vector for virus induced gene silencing in monocots and dicots. PLoS One 6, (2011).

87. Lück, S. et al. siRNA-Finder (si-Fi) Software for RNAi-Target Design and Off-Target Prediction. Front Plant Sci 10, (2019).

