## Supplemental Figures for "Long-read genome sequencing accelerated the cloning of *Pm69* by resolving the complexity of a rapidly evolving resistance gene cluster in wheat"

Li et al., 2022

**Supplementary Figures**

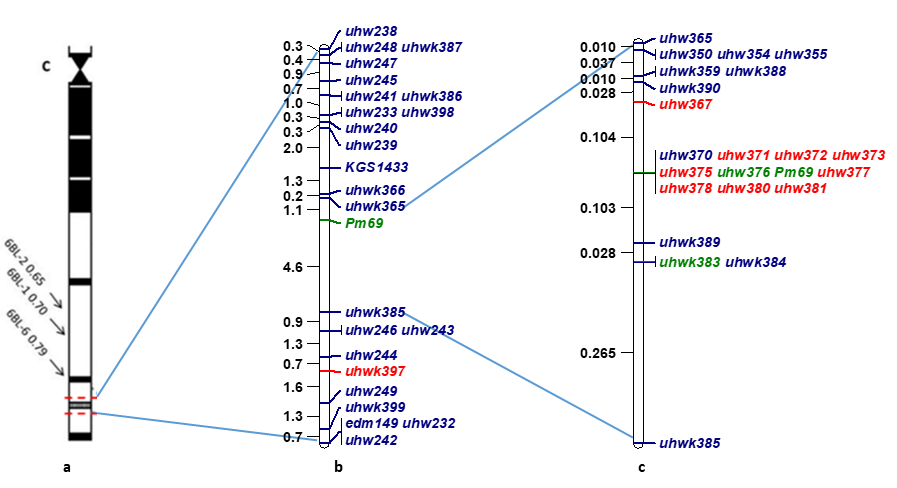

**Figure S1. The genetic map of *Pm69.* (a)** The deletion bin map of wheat chromosome arm 6BL; **(b)** Primary genetic map of the *Pm69* by using a small population (2500 F_2_ individuals). **(c)** High-resolution genetic map of *Pm69* on wheat chromosome 6BL. The mapping population of 147 homozygous recombinant inbred lines (RILs, F_6_ generation) which were developed from 5500 F_2_ individuals (G305-3M × LDN) based on the graphical genotyping approach. The blue markers, codominant; Red markers, LDN dominant; Green markers, G305-3M dominant.

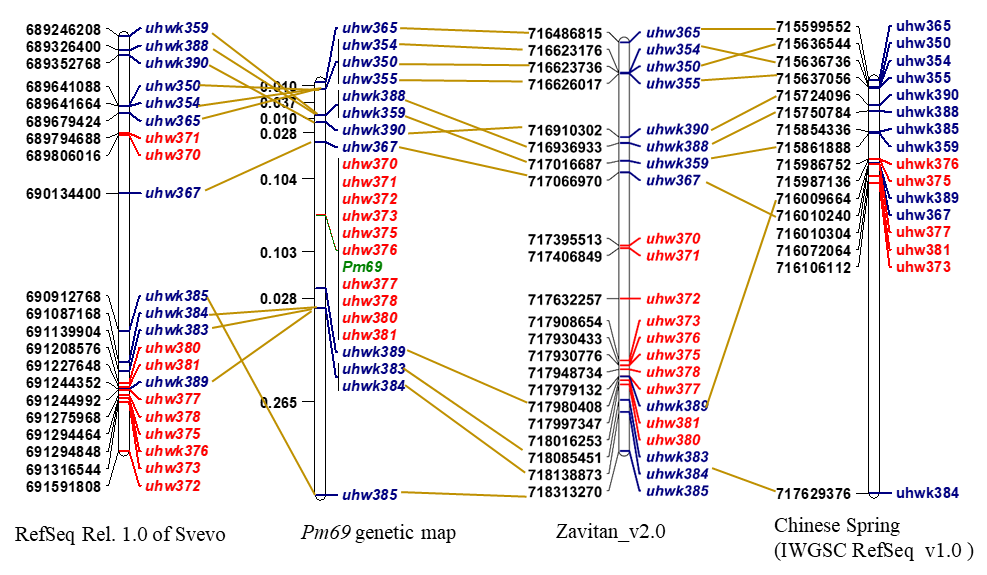

**Figure S2. Physical mapping of the *Pm69* region on chromosome 6BL based on the reference genomes of Zavitan, Svevo, and Chinese Spring (CS).** Anchoring of the sub-centiMorgan (cM) genetic map of the *Pm69* locus to the reference genomes of WEW_v2.0, durum wheat Svevo RefSeq Rel.1.0, and bread wheat CS IWGSC RefSeq v1.0. The red color is marking co-segregating markers to the *Pm69* phenotype. The blue color is indicating the flanking markers.

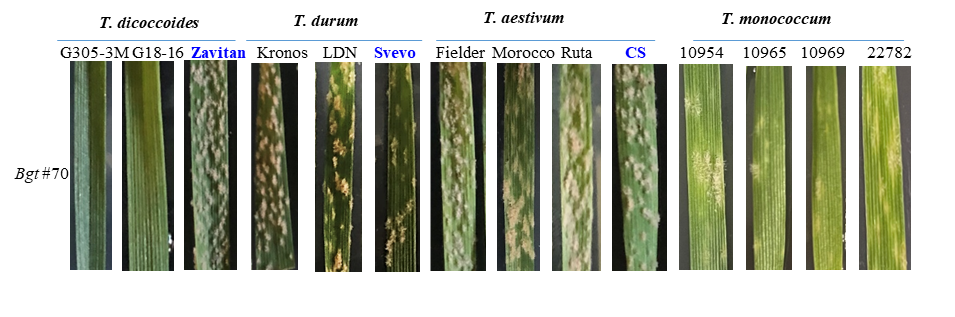

**Figure S3. The phenotypic response of different wheat accessions to *Bgt* #70.** *Bgt* #70 showed a strong virulence to several wheat species. WEW accession: Zavitan; Durum accessions: Kronos, LDN and Svevo; Bread wheat accessions: Fielder, Morocco, Ruta and Chinese Spring; *T. monococcum* accessions: NGB10953, NGB10965, NGB10969 and NGB22782, are susceptible to *Bgt* #70. The WEW accession: G305-3M and G18-16 that contained functional *Pm* genes (*Pm69* and *TdPm60*, respectively) showed high resistance to *Bgt* #70. The Zavitan, Svevo and CS were highly susceptible to *Bgt* #70, suggesting that the three reference genomes in Figure S2 did not contain the functional *Pm69* allele.

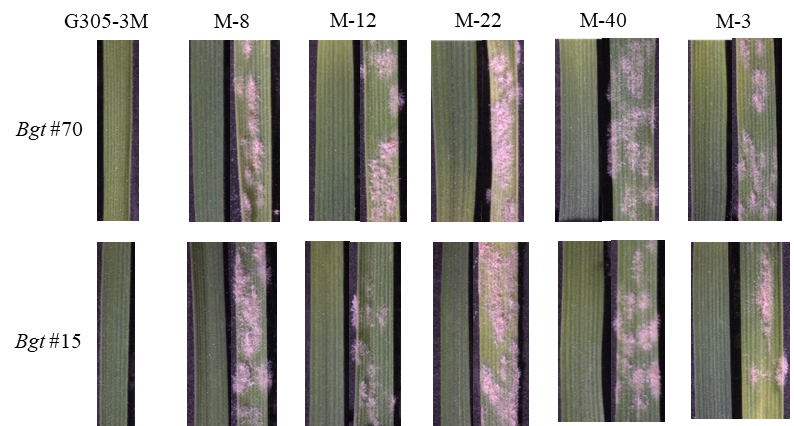

**Figure S4. The phenotypic response of EMS-derived mutants to *Bgt* #70 and *Bgt* #15**. The resistant plant is the sister line of each mutant, which was derived from the same M_0_ plant. *Bgt* #15 is another isolate was also used for *Pm69* genetic mapping in our previous report^1^.

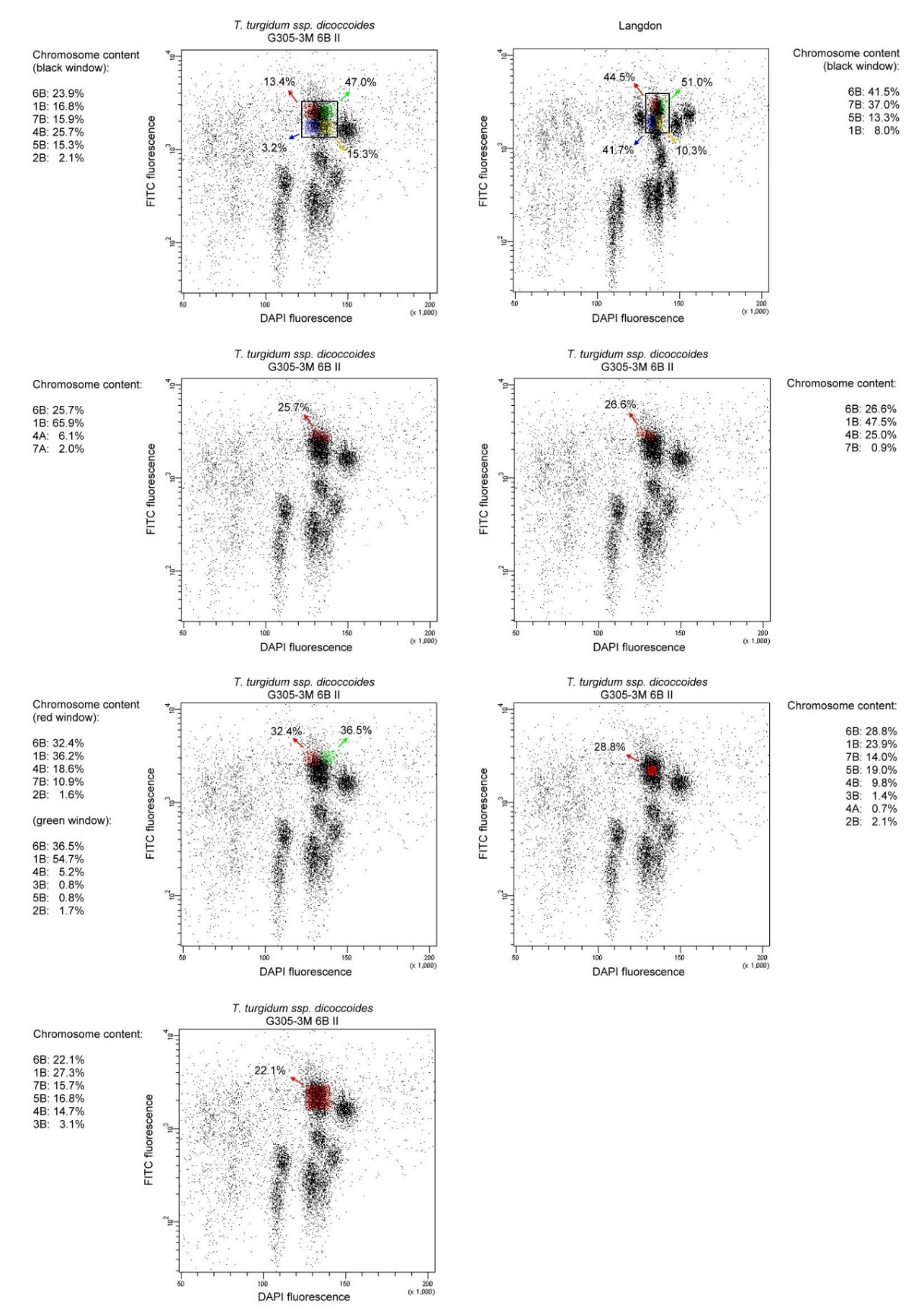

**Figure S5. The chromosome 6B sorting from G305-3M and LDN wheat lines.** The regions of bivariate (DAPI vs. FITC) flow karyotypes tested for 6B sorting are highlighted (black, red, and green) and their chromosome content in per cent obtained by microscopy investigations is given.

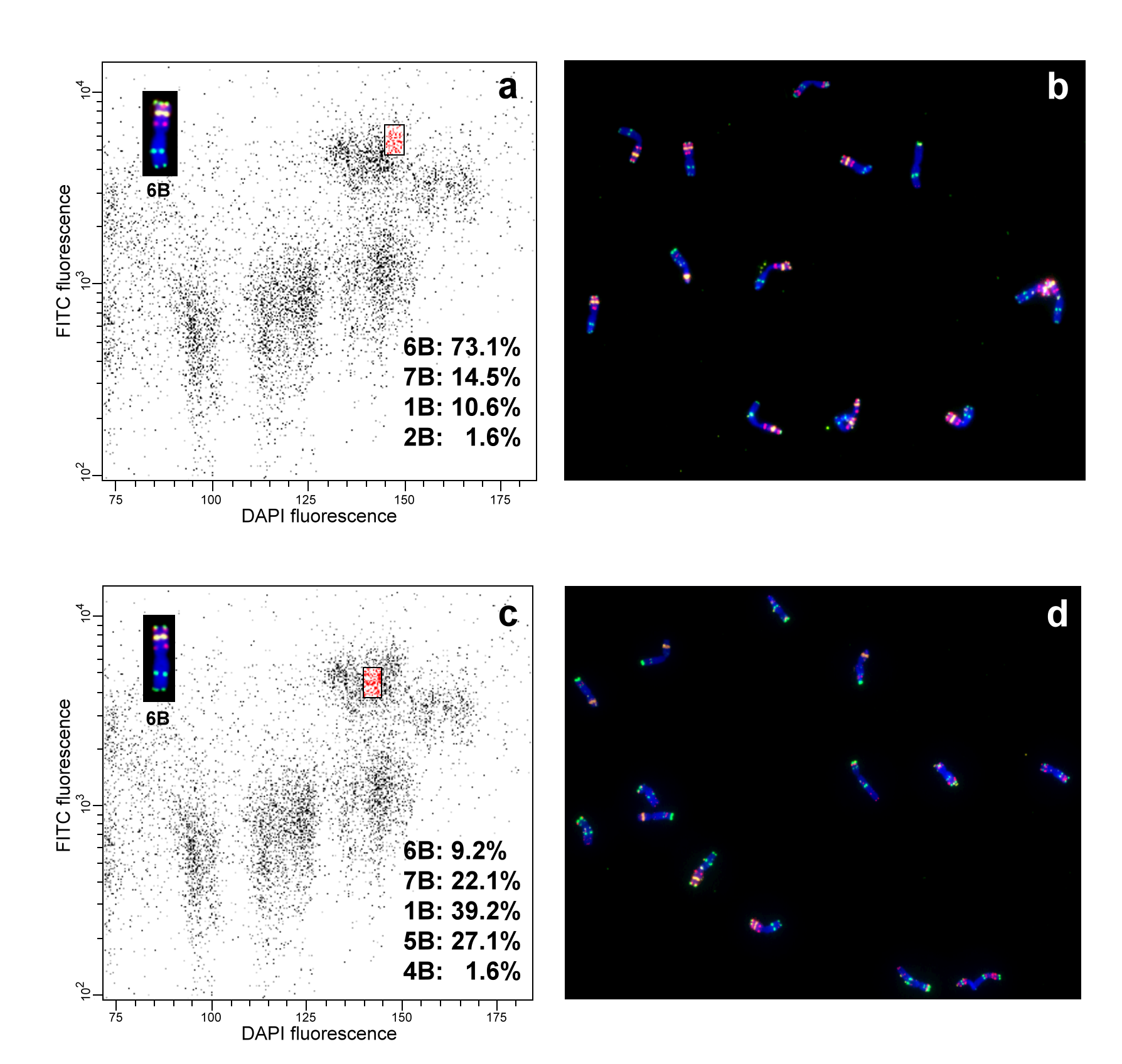

**Figure S6. Flow karyotyping and chromosome sorting from the introgression line SC28RRR-26.** Bivariate (DAPI vs. FITC) flow karyotype of the line SC28RRR-26 was obtained after the analysis of DAPI-stained chromosome suspension labelled by FISHIS using probes GAA-FITC and ACG-FITC. Two different positions of sort windows highlighted in red were tested to sort chromosome 6B (a and c). Chromosomes were flow-sorted from the colored regions onto microscope slides and identified by fluorescence *in situ* hybridization with probes for DNA repeats pSc119.2 (red), Afa family (green) and 18S rDNA (yellow) in different sorting windows (**b** and **d**).

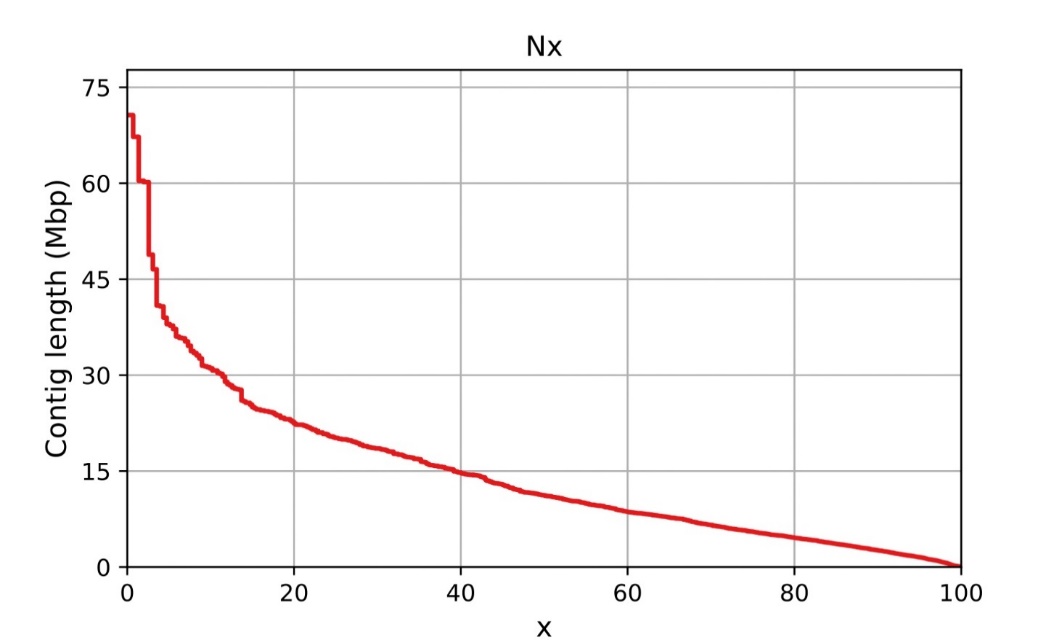

**Figure S7. Nx-length plot for ONT assembly of G305-3M.** Each Nx value represents the shortest contig length when summed with all larger contigs totaling X% of the total assembly size.

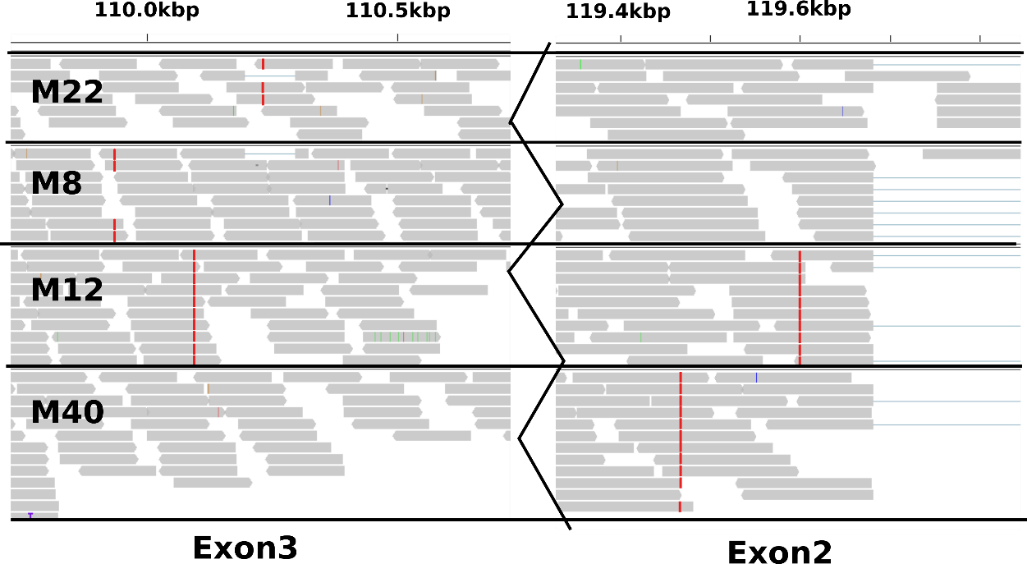

**Figure S8. The SNP information among the susceptible mutants viewed by IGV software in the exons of *NLR-6* gene.** The reads of susceptible mutants were aligned on the G305-3M ONT contig utg17163. The red vertical lines represent high-quality SNPs compared to the utg17163. The susceptible mutants were M-8, M-12, M-22, and M-40.

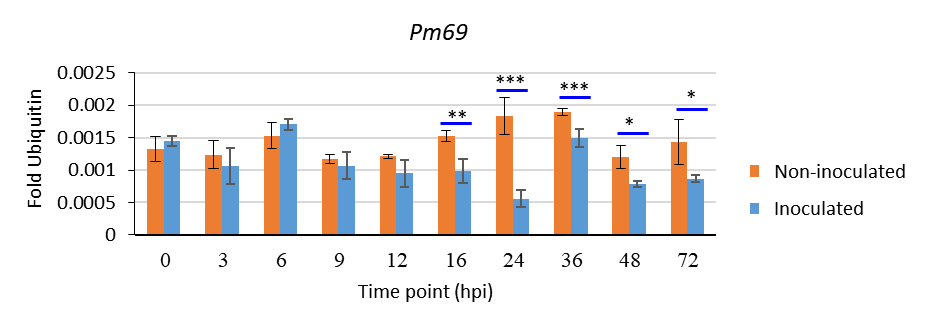

**Figure S9.** **The expression patterns of *Pm69* in G305-3M at the different *Bgt*-infection time points in both non-inoculated (mock) and inoculated plants.** The time points were ranged 0-72 hpi with *Bgt* #70. Asterisks indicate the level of significance by t-test of the *Pm69* expression (fold ubiquitin) in the target plants compared with the negative control of non-inoculated plant at each time point, p < 0.05 (*), p < 0.01 (**), p < 0.001 (***).

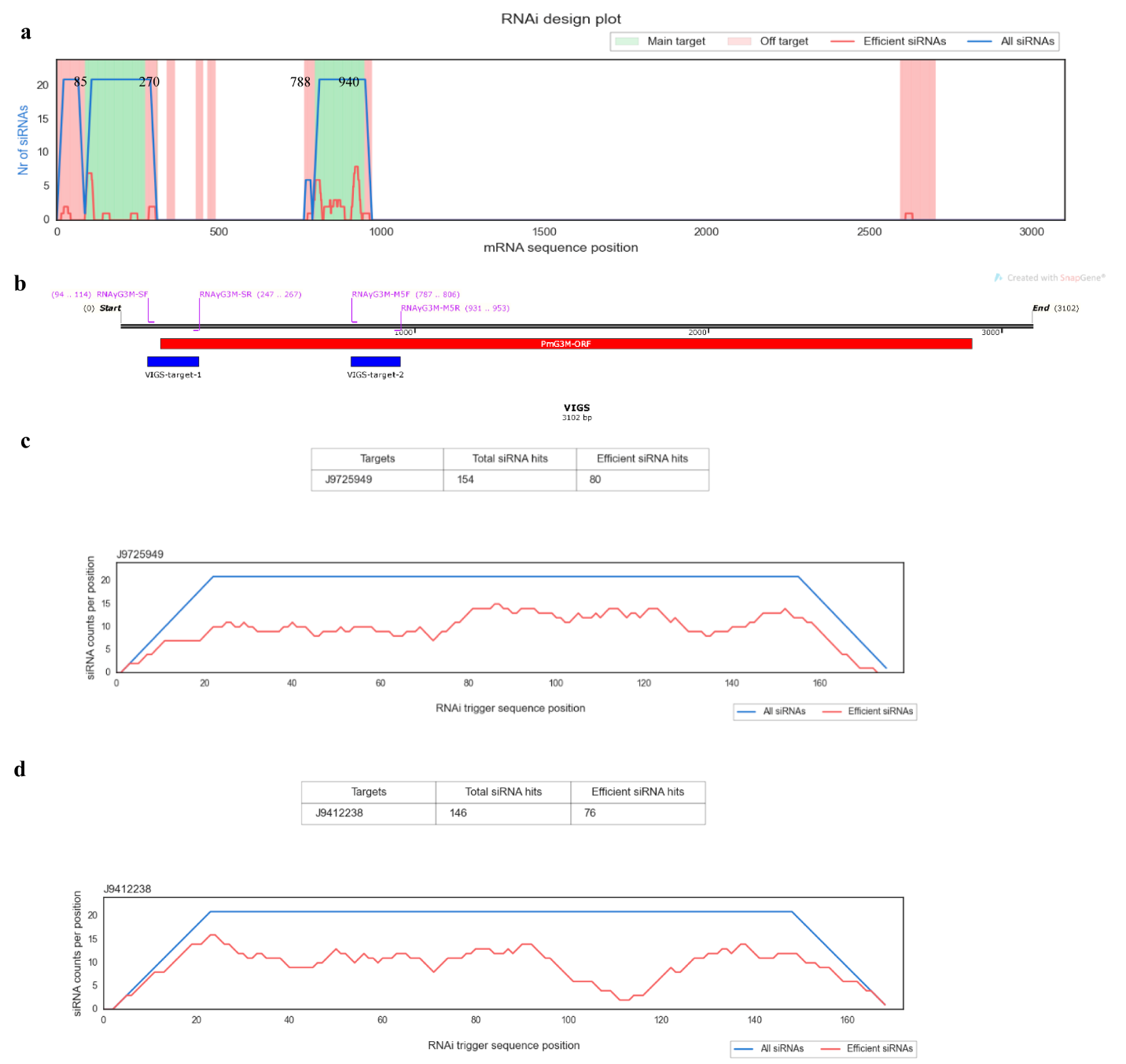

**Figure S10. The virus-induced gene silencing (VIGS) target region for silencing *Pm69*. (a)** Two specific regions were predicted by using si‐Fi software ^2^. The green locations are the predicted regions with efficient and less off-target siRNA among the G305-3M transcriptomes. **(b)** The selected VIGS target (blue bars) regions and two locations of primer pairs which used to amplify the target silencing region of *Pm69*. **(c)** The predicted off-target gene of VIGS-target1 (for BSMV:Pm69-1) sequence. **(d)** The predicted off-target gene of VIGS-target2 (for BSMV:Pm69-2) sequence. The two off-target genes were predicted as truncated genes in the G305-3M transcriptomes, which probably did not have the resistant function. Therefore, these two target parts could be used for *Pm69* silencing in VIGS system.

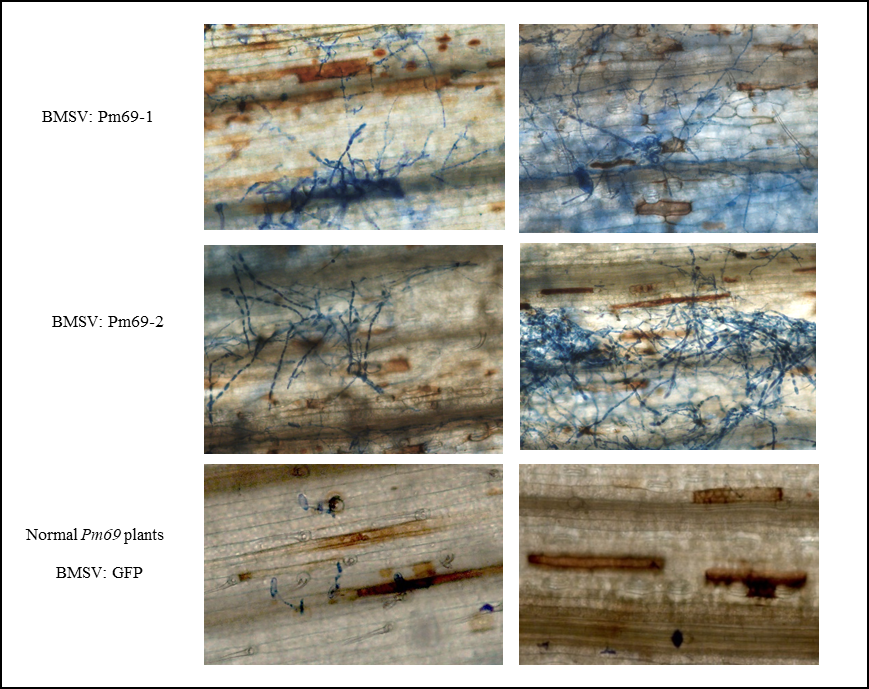

**Figure S11. The microscopy of ROS accumulation (reddish-brown coloration) and the fungal growth (blue-colored hyphae) in the *Pm69* wild-type and *Pm69*-silencing plants.** DAB and Coomassie blue staining of *Pm69* or GFP silencing leaves of introgression line SC28RRR-26 (Ruta +*Pm69*) at 7 dpi with *Bgt* #70. Construct of BMSV:GFP was used as the negative control for silencing the GFP genes. Constructs of BMSV:Pm69-1 and BMSV:Pm69-2 were used for silencing the *Pm69*.

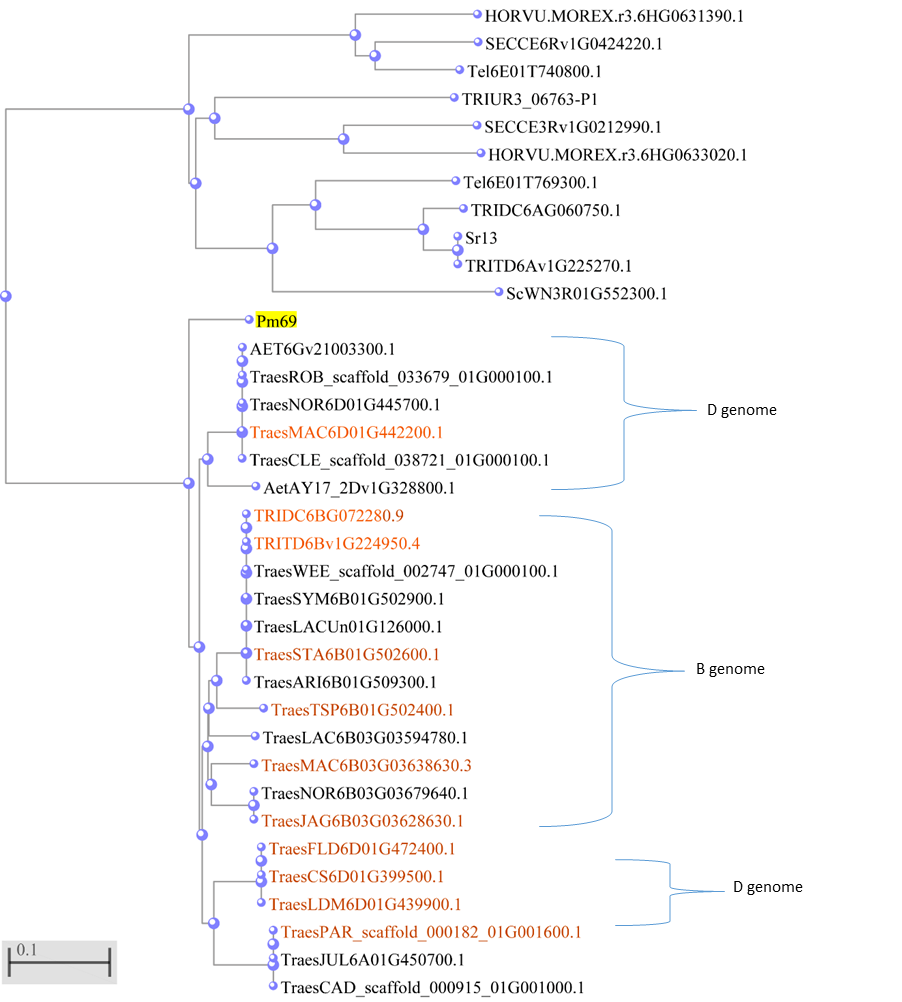

**Figure S12. Distance tree of Pm69 protein and its homologs** **or similar proteins.** Protein sequences can be found by using the ID names (present in the tree) from the website of WheatOmics 1.0, EnsemblPlants, or NCBI. The ID of Sr13 in GenBank: QQP18374.1. The orange color genes have been demonstrated as susceptible genotypes to *Bgt* #70, and the other alleles in the same branch suggested that they are identical alleles. Therefore, these identified *Pm69* homologs are nonfunctional alleles to *Bgt* #70.

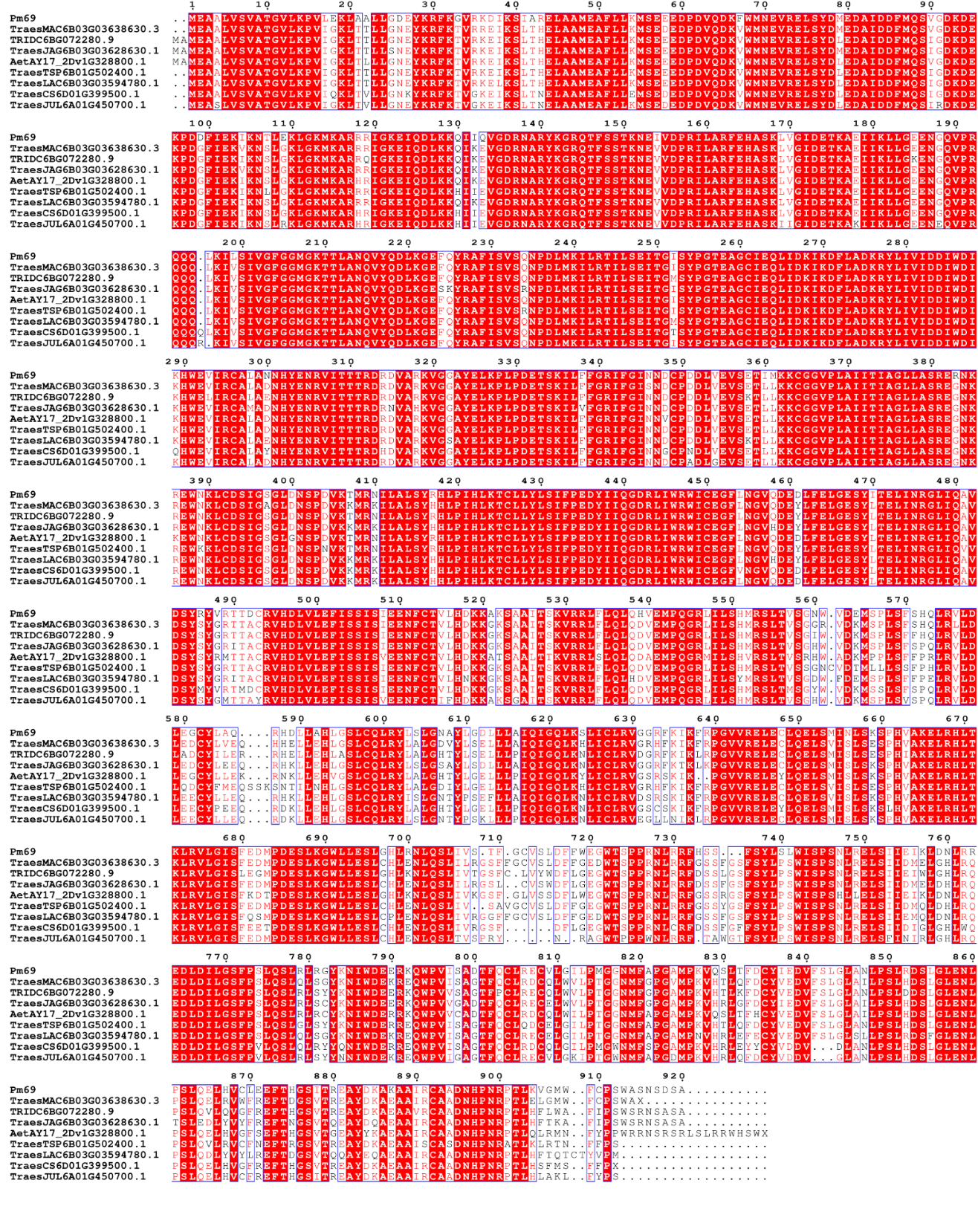

**Figure S13. Sequence alignments of Pm69 protein and eight representative *Pm69* homologs**. The red colored region presents the conserved part of the protein. The sequence of those homologs could be found on the website of WheatOmics 1.0 and EnsemblPlants by using their ID names. The alignments were constructed by the multiple alignment tool CLUSTALW from the NCBI and visualized by using a website tool (<https://espript.ibcp.fr/ESPript/cgi-bin/ESPript.cgi>).

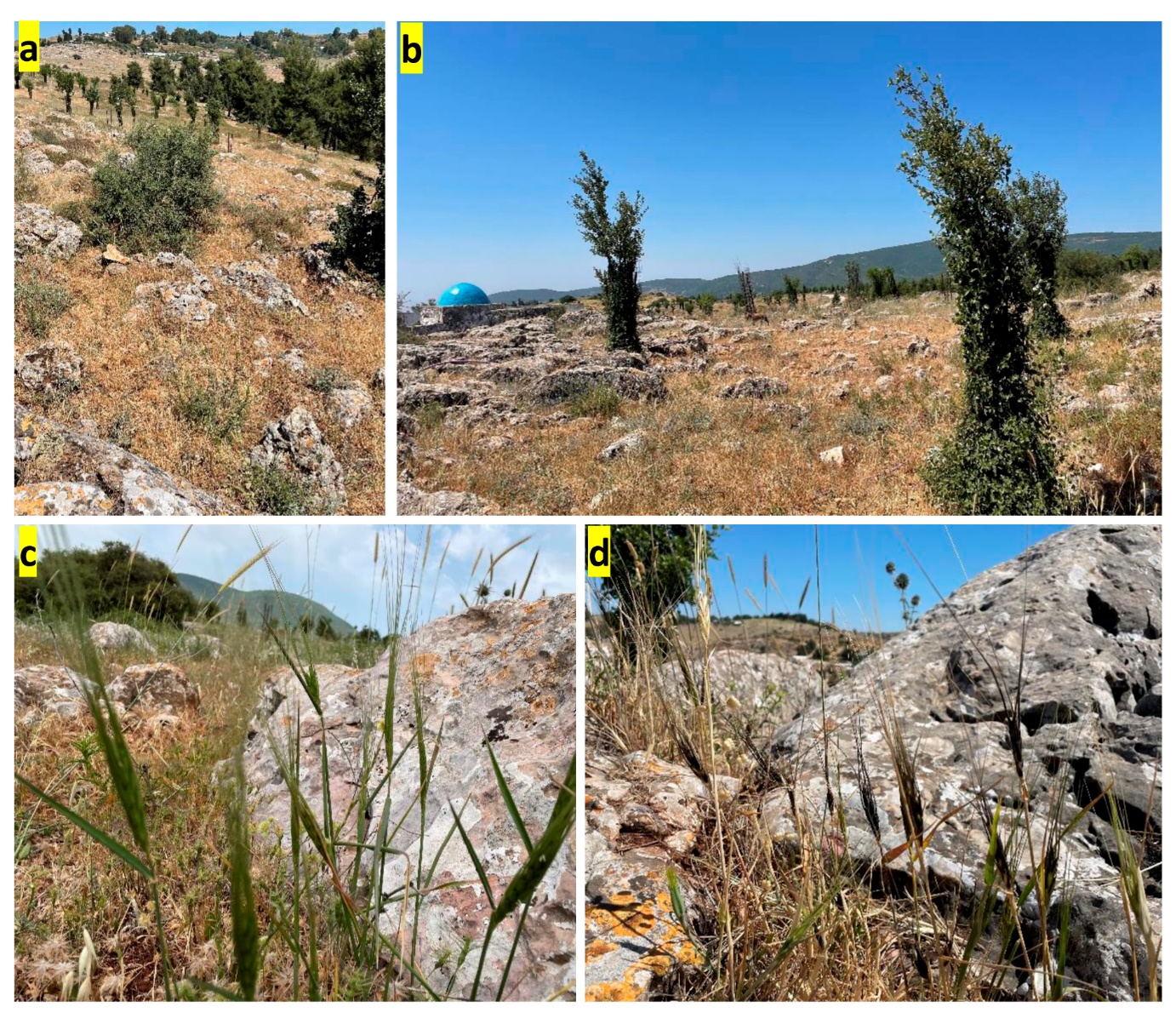

**Figure S14.** **The wild emmer wheat in the natural field.** **a** and **b**: around the *Pm69* lines collection site (south of Kadita, Northern Israel). **c**: WEW in the flowering period (09 May 2022). **d**: WEW in the maturation period (25 May 2022). WEW could only be found near the rocks in the natural field.

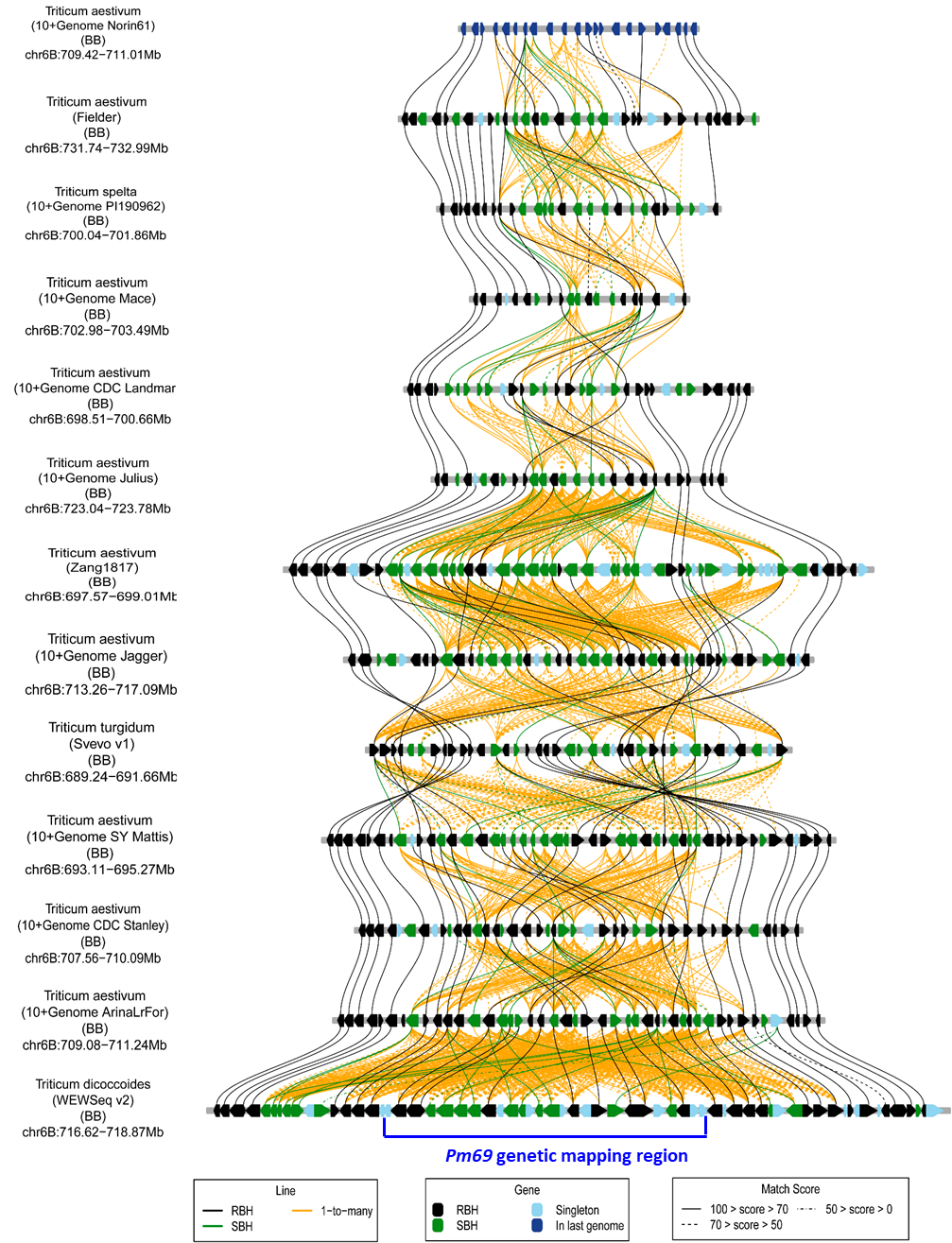

**Figure S15. Collinearity analysis of *Pm69* genetic region among different wheat genomes.** RBH: Reciprocal best hit; SBH: single-side best hit. The Collinearity was analyzed by the website of <http://wheat.cau.edu.cn/TGT/zavitanv2_TGT/?navbar=MicroCollinearity>. Access date: 03 February 2022. Chinese Spring IWGSC RefSeqv1.1 was not included here because of the inversion structure, probably by the assembly mistakes, which have been shown in Figure 6 and Table S15. The Norin61 was not included because of the lower collinearity, in which just two genes were present the collinearity and shown in Table S15. The yellow lines showed the genes target more than one matched gene, probably because of the NLR duplication events.

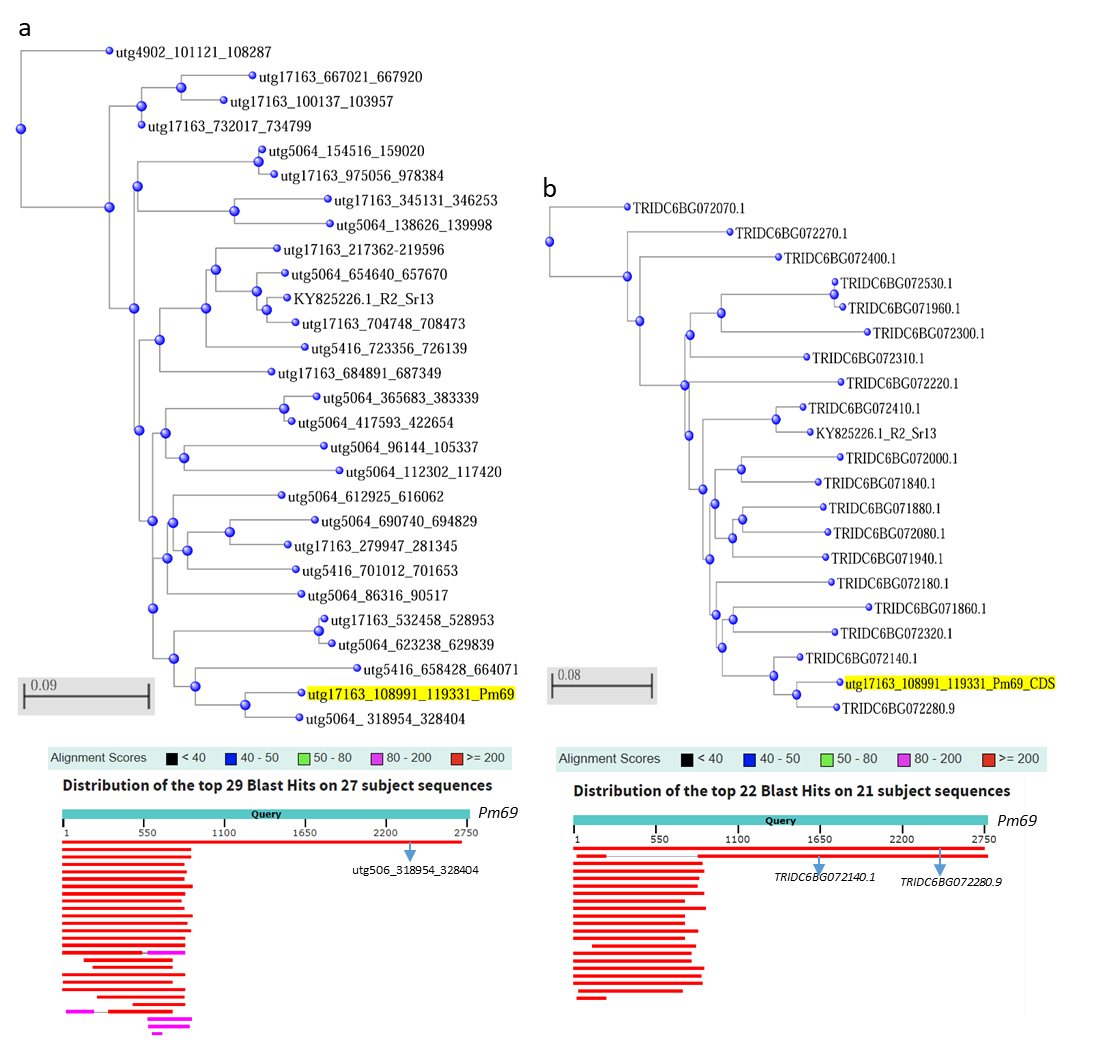

**Figure S16. Distance trees of NLRs in the *Pm69*/*pm69* NLR clusters of G305-3M (a) and Zavitan (b).** *Sr13* is a control for the genetic distance of *Pm69*. The CDS sequences of the genes were analyzed in the NCBI BLASTN programs tools by using the *Pm69* sequence as a probe for the blast. The Rx_N (1-291) and NB-ARC (516-1248) were more conserved than the LRR (1647-2124) domain by comparing *Pm69* with these *NLRs* from the NLR cluster.

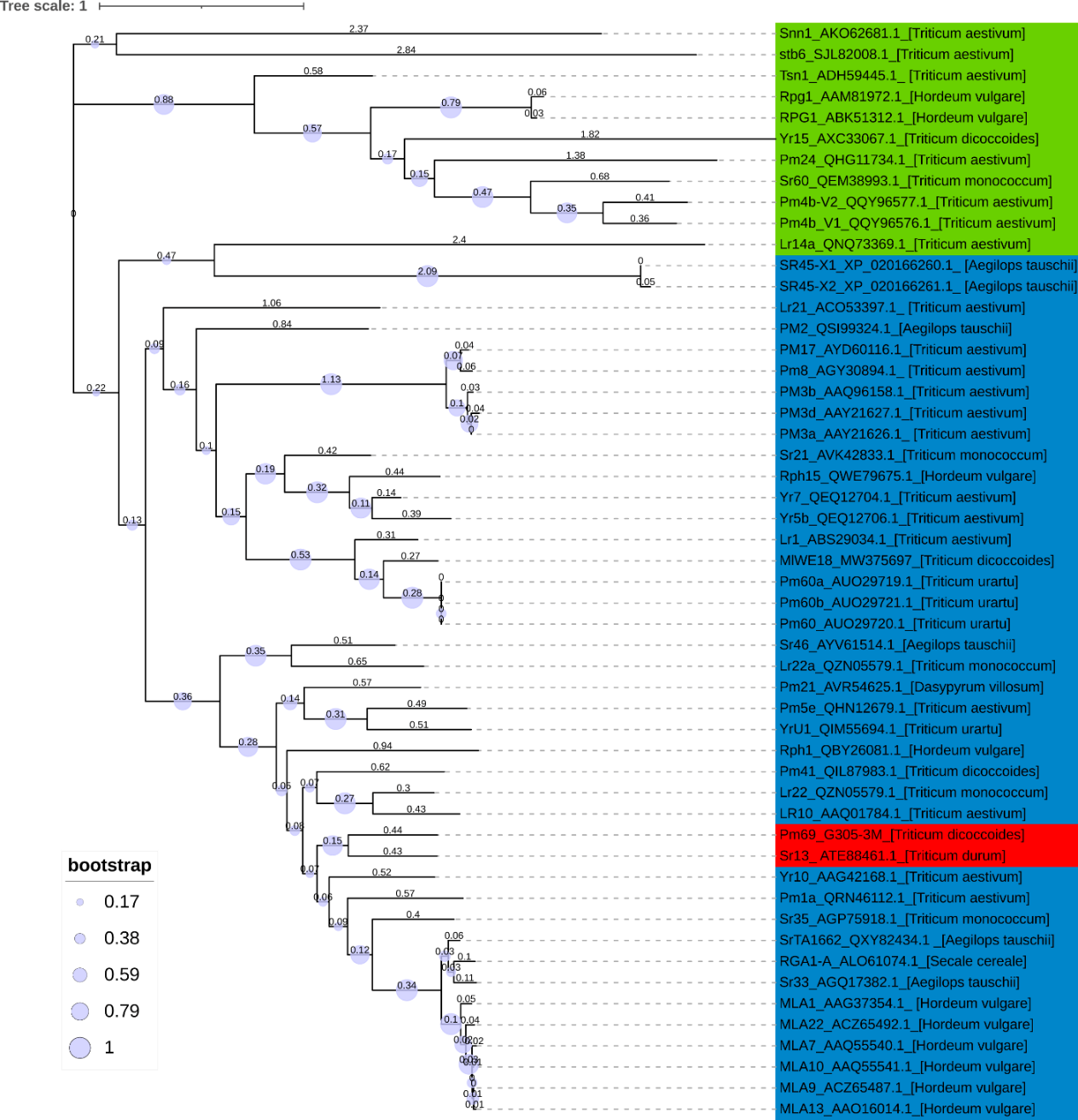

**Figure S17. Phylogenetic tree of Pm69 and 36 published resistance proteins.** The protein sequence could be found by the ID numbers in the NCBI database. The blue-colored protein contains the NLR structure. The green-colored protein contains are nonconventional R proteins without NLR structures. The red-colored proteins are Pm69 and Sr13. The evolution tree is built by MEGA-X (aligning by MUSCLE, Neighbor-Joining algorithm, bootstrap: 500 times) and visualized on the website of iTOL v6 (<https://itol.embl.de/>). The circle area represents the number of the bootstrap. The horizontal lines are branches and represent evolutionary changes (above numbers) measured in a unit of time or genetic divergence.

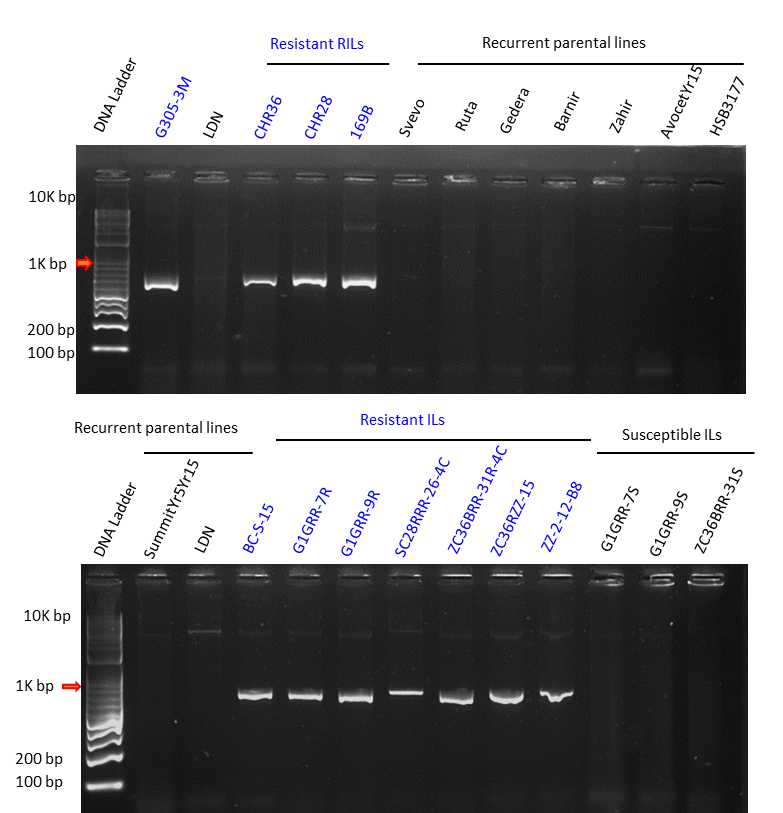

**Figure S18. Representative 1% agarose gel electrophoresis of PCR products amplified by *uhw430* for the selection of introgression lines (ILs)*.*** The resistant wheat RILs and ILs (blue colored) showed positive PCR amplification of *uhw430.*

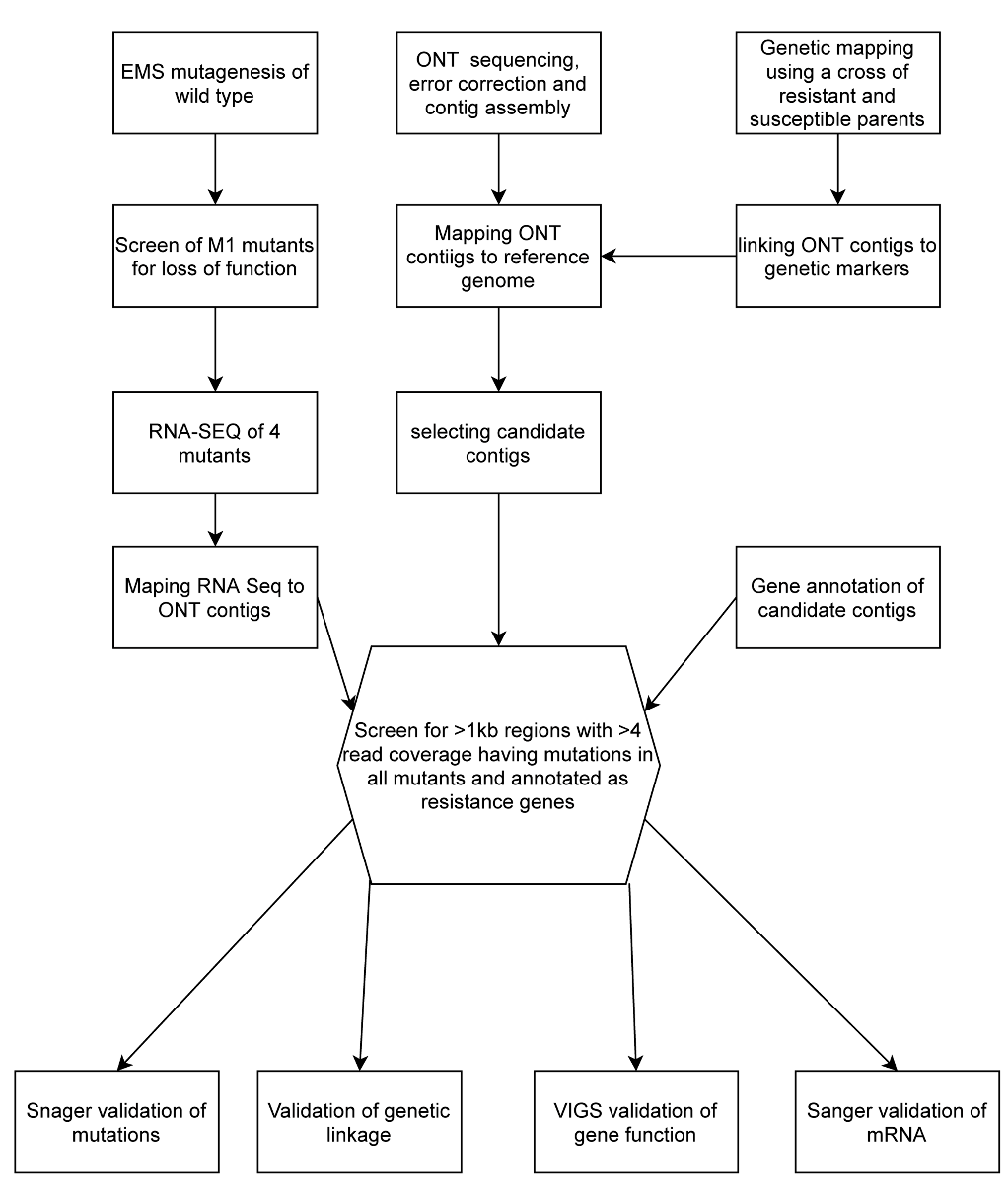

**Figure S19. The workflow of gene cloning by** **ONT-MutRNAseq.**

**References**

1. Xie, W. et al. Identification and characterization of a novel powdery mildew resistance gene *PmG3M* derived from wild emmer wheat, *Triticum dicoccoides*. *Theoretical and Applied Genetics* 124, 911–922 (2012).
2. Lück, S. et al. siRNA-Finder (si-Fi) software for RNAi-target design and off-target prediction. *Frontiers in Plant Science* 10, (2019).
